# SRS microscopy identifies inhibition of vitellogenesis as a mediator of lifespan extension by caloric restriction in *C. elegans*

**DOI:** 10.1101/2025.01.31.636008

**Authors:** Bowen Yang, Bryce Manifold, Wuji Han, Catherin DeSousa, Wanyi Zhu, Aaron Streets, Denis V. Titov

## Abstract

The molecular mechanisms of aging are not fully understood. Here, we used label-free Stimulated Raman scattering (SRS) microscopy to investigate changes in proteins and lipids throughout the lifespan of *C. elegans*. We observed a dramatic buildup of proteins within the body cavity or pseudocoelom of aged adults that was blunted by interventions that extend lifespan: caloric restriction (CR) and the reduced insulin/insulin-like growth factor signaling (IIS) pathway. Using a combination of microscopy, proteomic analysis, and validation with mutant strains, we identified vitellogenins as the key molecular components of the protein buildup in the pseudocoelom. Vitellogenins shuttle nutrients from intestine to embryos and are homologous to human apolipoprotein B, the causal driver of cardiovascular disease. We then showed that CR and knockdown of vitellogenins both extend lifespan by >60%, but their combination has no additional effect on lifespan, suggesting that CR extends the lifespan of *C. elegans* in part by inhibiting vitellogenesis. The extensive dataset of more than 12,000 images stitched into over 350 whole-animal SRS images of *C. elegans* at different ages and subjected to different longevity intervention will be a valuable resource for researchers interested in aging.

## INTRODUCTION

Aging is a nearly universal process, yet the molecular mechanisms underlying this process are not fully understood^1^ . The nematode *C. elegans* is one of the most common model organisms used to study the aging process because of its short lifespan and well-characterized biology. Multiple genetic and dietary interventions, including reduced insulin/insulin-like growth factor signaling (IIS), caloric restriction (CR), and respiratory inhibition, have been shown to extend lifespan across various model organisms, including *C. elegans*^2–5^. However, the downstream players of these pro-longevity interventions and the extent to which they operate through distinct or overlapping mechanisms have yet to be fully elucidated.

Vitellogenins are a class of proteins that have been linked to aging in *C. elegans*. In young adults, these proteins function to transport amino acids and lipids from the intestine to the germline to support oocyte development^6^. Six vitellogenin genes expressed in the intestine produce four yolk protein polypeptides–YP170A, YP170B, YP115, and YP88^7,8^. With age, vitellogenin protein levels increase dramatically, while pro-longevity interventions like CR and reduced IIS are associated with their decreased expression^6,9,10^. Notably, reducing vitellogenin levels has been shown to extend lifespan in *C. elegans*^9–13^. Vitellogenins share functional and structural similarities with human apolipoprotein B (ApoB), including sequence homology, nutrient transport function, and receptor mediated endocytosis in target tissues. ApoB is a key structural component of atherogenic lipoproteins including LDL, which transports cholesterol and triglycerides in the bloodstream^14^. Elevated ApoB levels are a major driver of cardiovascular disease, the leading cause of death in the United States^15^. Like vitellogenins, ApoB levels in humans increase with age and are modulated by dietary changes^14,16–18^.

In this work we perform stimulated Raman scattering (SRS) microscopy on whole live *C. elegans* during aging and a variety of longevity interventions. SRS microscopy is a technique that provides visual contrast based on molecular vibrations inherent to endogenous biomolecules in a sample. SRS microscopy has been previously shown as a valuable tool specifically in the study of *C. elegans* given its power in observing metabolic changes in worms, especially regarding lipogenesis. In fact, some of the first studies introducing SRS microscopy as a tool for bioimaging demonstrate imaging of *C. elegans*^19–21^. Beyond these early demonstrations, SRS microscopy has contributed extensively to understanding lipid droplet biology and metabolism^22–25^, cholesterol presence^26,27^, and retinoid storage in *C. elegans*^28,29^. Further, deuterium isotope tracing in SRS has recently been leveraged to understand the spatiotemporal dynamics of metabolism in worms and in aging drosophila^30–33^. While these previous works have demonstrated many interesting findings in worms, they have largely been focused on the imaging of gut lipids and lipogenesis via SRS, often at a single Raman transition. However, systematic characterization of whole-animals throughout lifespan and in response lifespan interventions has not been done.

Here, we harness SRS microscopy to characterize spatiotemporal changes in both proteins and lipids across the entire *C. elegans* body and throughout lifespan under two longevity interventions: CR and reduced IIS. Through this work we have produced 355 stitched whole-animal SRS images that are compiled in an accessible database for the use of the aging community, along with information such as age, strain, and diet. Our extensive imaging dataset revealed significant protein buildup in the pseudocoelomic region with age that was prevented by CR and reduced IIS. Through analysis of proteomics datasets and composite 2 photon fluorescence (2PF) microscopy, we found that a class of proteins called vitellogenins form the major component of this protein buildup. We then demonstrated that inhibition of vitellogenesis and CR both extend lifespan by ∼60% and that the lifespan of CR animals was not further extended by inhibition of vitellogenesis. Furthermore, we showed that known mutants that prevent CR-mediated lifespan extension are regulators of vitellogenesis. Our results support a hypothesis that CR extends the lifespan of *C. elegans* in part by inhibiting vitellogenesis.

## RESULTS

### Stimulated Raman scattering microscopy identifies extensive protein buildup in the pseudocoelomic space of aged C. elegans

An example of *C. elegans* imaged in two color SRS is shown in Figures 1A-1F. Here the two Raman transitions are acquired and then composited into one image. Specifically, the two Raman transitions are 2850 cm^-1^ (shown in gold) and 2920 cm^-1^ (shown in blue). These two transitions are canonical choices in SRS imaging of the carbon-hydrogen (CH) region of the Raman spectrum (2800-3100 cm^-1^) corresponding to CH2 and CH3 stretches respectively. The CH2 (2850 cm^-1^, gold) peak in unlabeled samples is generated mostly by lipid molecules and is thus referred to commonly as the lipid peak. The CH3 (2920 cm^-1^, blue) peak, meanwhile, is generated in part by lipid molecules, but most dominantly by proteins in cells. Being that proteins and lipids make up most of the dry mass of cells, these two peaks are commonly chosen to visualize cells and tissues in label-free SRS imaging. The imaging parameters (e.g., laser powers, objective, gains, etc.) were held constant across all images acquired in this study, so the relative pixel intensities or brightnesses of gold/blue pixels can be compared across images and represent relative concentrations of Raman active molecules.

**Figure 1.**
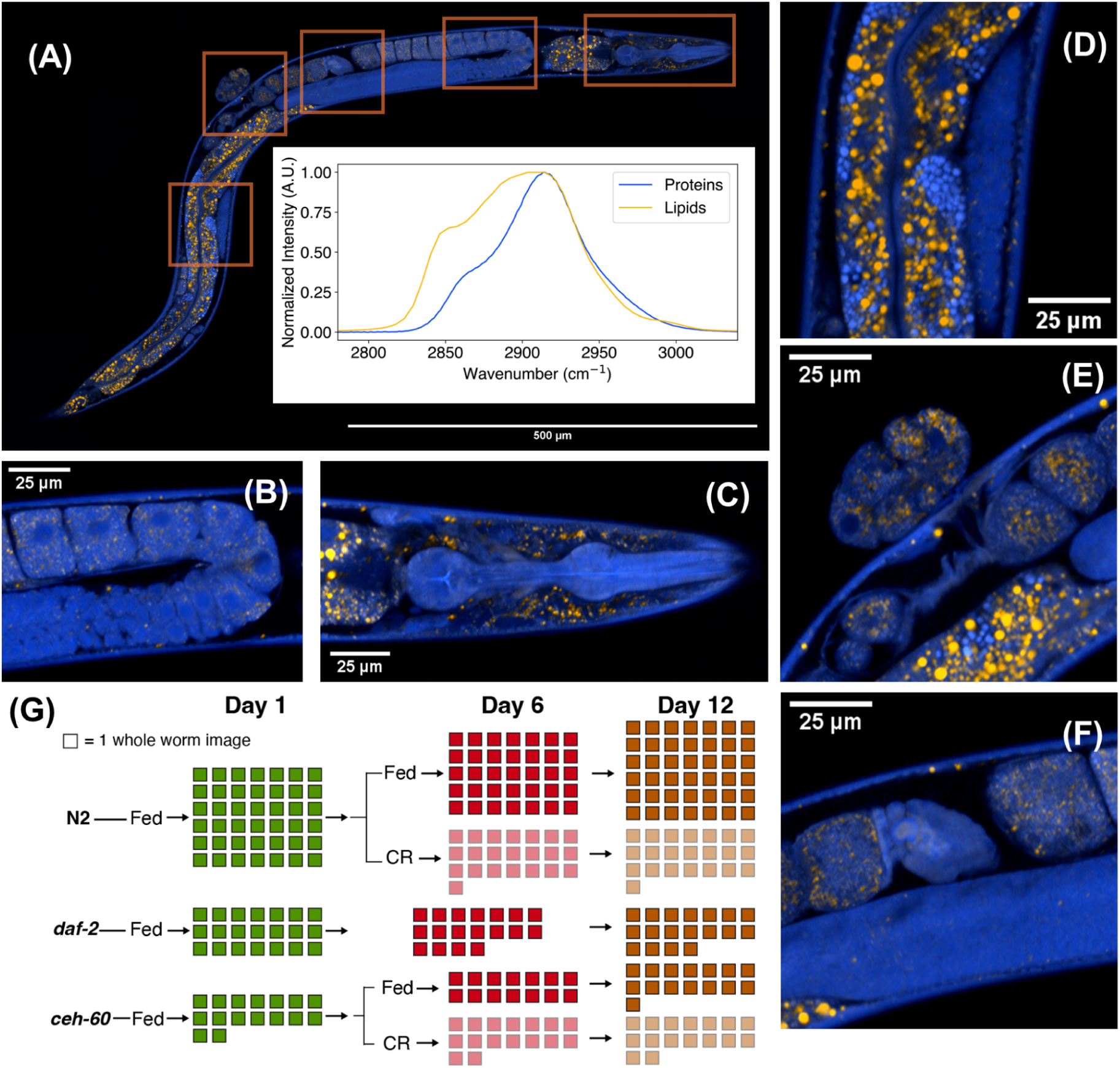
Whole-body SRS images of adult *C. elegans* **(A)** Whole-body SRS image of an N2 (wildtype) Day 1 Adult worm. Blue represents protein. Gold represents lipids. Orange boxes highlight the inset zoom-ins of the subsequent panels. The inset graph shows the respective normalized pure spectra of proteins and lipids highlighting the CH2 peak (∼2850 cm^-1^) and CH3 peak (∼2920 cm^-1^). **(B)** Inset image from (A) of the gonad. Here the progressive activation and growth of oocytes is evident. **(C)** Inset image from (A) of the head. The bulb, pharynx, and anterior intestine are visible. The nerve ring structures are also seen. **(D)** Inset image from (A) of the intestine. Here the main tract and lumen are well defined, with both lipid droplets (gold) and gut granules (blue) seen. A fold of the posterior gonad is also present. **(E)** Inset image from (A) showing a recently laid egg. The vulva and internally-developing eggs are also visible. **(F)** Inset image from (A) of the spermatheca. Diagram outlining SRS images taken across strains, conditions, and age where each box represents a one whole worm image.

For the 355 worms imaged in this study (Figure 1G), multiple fields of view were acquired in a grid around the worm and subsequently stitched together to create whole worm images. Within the example worm shown in Figures 1A-1F, many of the expected internal somatic features and organelles are clearly observable, all without the need for exogenous labeling here. For example, the anterior mouth, pharynx, and nerve ring; the gonadal track with developing oocytes, spermatheca, and newly laid egg; and intestinal tract including both the expected lipid droplets and lysosomal-related gut granules. Here, these features can be easily identified and compared amongst ages and intervention conditions to understand *C. elegans* aging quantitatively and holistically, without the bias of labeling specific markers.

To investigate age-related changes in the spatial distribution and concentration of proteins and lipids in *C. elegans* age, we performed SRS imaging at days 1, 6, and 12 of adulthood (Figure 2A-C). These time points represent the early, middle, and late stages of the lifespan in *ad libitum* fed animals, as determined by our lifespan measurements (Figure 3F).

**Figure 2.**
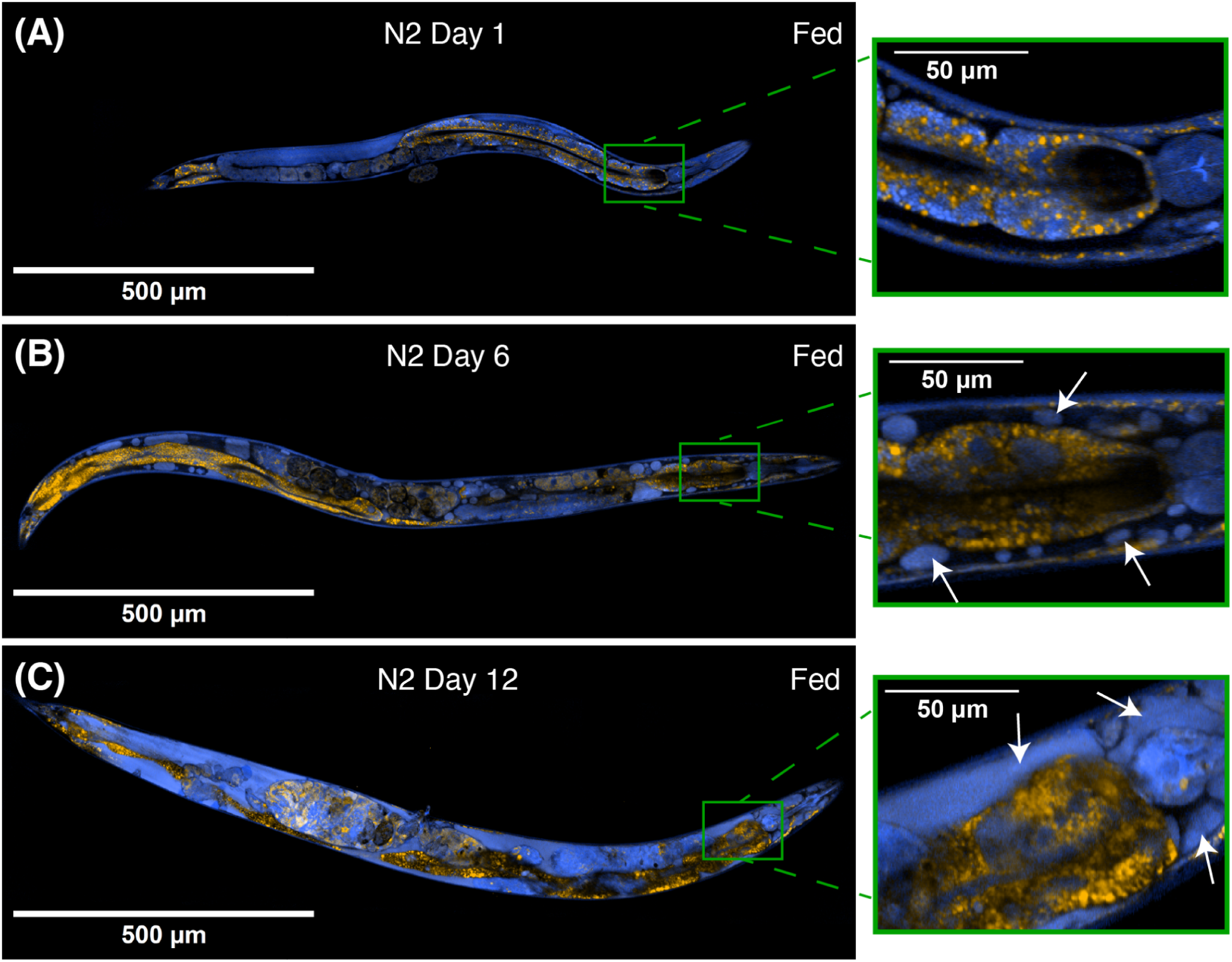
SRS microscopy reveals extensive protein buildup in pseudocoelomic space of aged worms **(A)** SRS image of N2 Day 1 Adult and inset of the pharynx area. **(B)** SRS image of N2 Day 6 Adult and inset showing the increase in lipid droplets in the intestine and beginning features of protein build-up in the pseudocoelom labelled with white arrows. SRS image of N2 Day 12 Adult and inset showing the extensive build-up of protein in the pseudocoelom labelled with white arrows.

**Figure 3.**
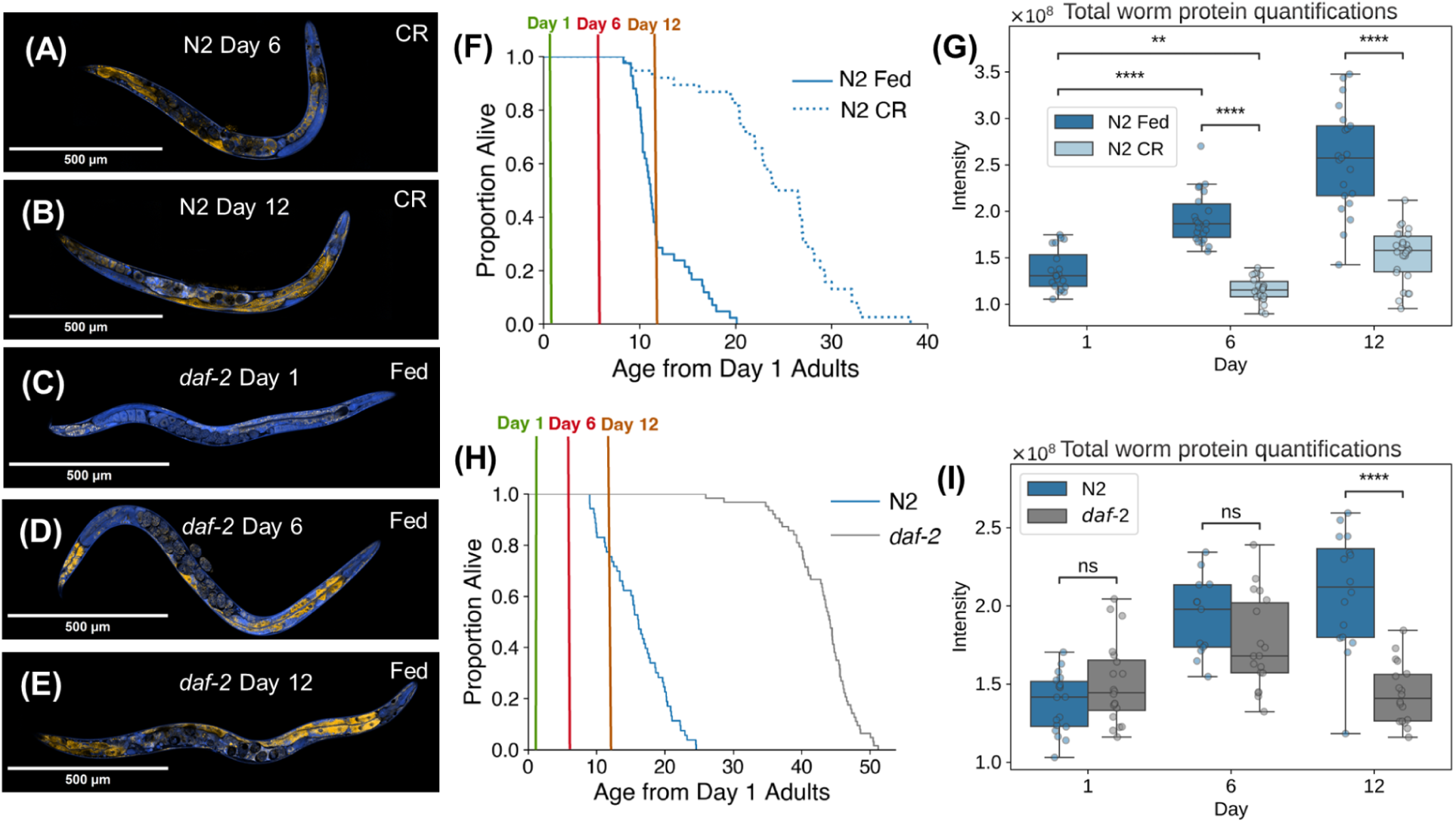
Caloric restriction and reduced insulin/insulin-like growth factor signaling mitigate age-related increase in protein buildup **(A)** SRS image of N2 Day 6 Adult, CR **(B)** SRS image of N2 Day 12 Adult, CR **(C)** SRS image of *daf-2* mutant Day 1 Adult, fed **(D)** SRS image of *daf-2* mutant Day 6 Adult, fed **(E)** SRS image of *daf-2* mutant Day 12 Adult, fed **(F)** Lifespan of N2 fed and CR with imaging time points highlighted **(G)** Quantification of total protein for N2 fed and CR. **(H)** Lifespan of N2 and *daf-2* mutant with imaging time points highlighted **(I)** Quantification of total protein for N2 and *daf-2* mutants *, **, ***, and **** indicate p < 0.05, 0.01, 0.001, and 0.0001 (Welch’s t-test), respectively. Horizontal bars of the boxes represent maximum, 0.75, 0.5, 0.25, minimum, respectively. Outliers are determined by whether the point deviates 1.5 times the interquartile range (0.75 quartile - 0.25 quartile), from either quartile.

SRS imaging reveals several previously reported features of young and aged worms. As expected, lipid signals were predominantly localized in the intestine, consistent with prior studies identifying this tissue as the primary site of lipid storage in *C. elegans* (Figure 1A and 1D)^21,34–36^. Additionally, in day 12 adults, we observed two prominent “masses” enriched in both protein and lipid within the gonadal region (Figure 2C and S5). These closely resemble the previously reported “uterine tumours” observed in old worms^37^.

Our SRS images also uncovered new insights into protein and lipid distribution. As worms aged, tissues became increasingly disorganized, with an apparent loss of structural integrity between the tissues (Figure 2B and 2C). Most notably, we observed a pronounced buildup of proteins within the pseudocoelomic region as worms aged (Figure 2B and 2C). By day 12, this protein buildup was dramatic, with many of the worms displaying extensive protein enrichment throughout their body cavities.

### Caloric restriction and reduced insulin/insulin-like growth factor signaling mitigate age-related increase in protein buildup

We tested whether established lifespan-extending interventions–CR and reduced IIS–could mitigate the age-related protein buildup. To induce CR, we used a solid dietary restriction protocol starting on day 1 of adulthood, which extends lifespan by >90%^4^ (Figure 3F). As previously reported, CR reduced lipid levels and body size compared to *ad libitum* fed animals (Figure S1)^5,6^. Notably, CR worms displayed significantly reduced levels of protein buildup compared to fed animals (Figure 2B, Figure 2C, Figure 3A, 3B, and 3G, Figure S1). Analysis of many independent worms shows that the attenuation of protein buildup is even observed as a decrease in total and average protein signal (Figure 3G, Figure S1).

We next investigated the effects of the reduced IIS on the age-related protein buildup using the *daf-2* (e1370) allele^3,11,38^. Since this temperature-sensitive mutant requires lower temperatures during development to prevent dauer formation, we included corresponding wildtype N2 controls grown under identical conditions (Figure S5). We first confirmed that these *daf-2* mutant worms lived significantly longer compared to their wildtype counterparts (Figure 3H). Then, using these same conditions, we performed SRS imaging at day 1, day 6, and day 12 of adulthood. Consistent with previous studies, *daf-2* mutant worms displayed increased lipid levels in their intestine by day 6 of adulthood (Figures 2B, 3D, and S1)^21,36,39–42^. Importantly, similar to CR, *daf-2* worms showed significantly lower levels of protein buildup by day 12 of adulthood compared to their N2 counterparts (Figure 3D, 3E, and 3I).

### Proteome-wide analysis reveals that vitellogenins show the largest age-related increase in absolute abundance

To identify proteins that may contribute to the observed protein buildup with age, we applied a computational approach using published proteomics data^38,43^. Using whole-body, label-free proteomics data collected across the lifespan of *C. elegans*, we estimated the fraction each protein occupied relative to the total proteome at each time point. This approach, previously shown to accurately quantify protein levels across five orders of magnitude^44^, allowed us to determine changes in absolute protein composition across age. We then developed an algorithm to rank proteins based on their change in proteome fraction with age, from the lowest to highest (Figure S3). Our analysis revealed that most proteins showed minimal changes in their fraction of total proteome occupied with age (Figure 4A and 4B). Across the two independent datasets only 0.85-1.1% of identified proteins exhibited an increase of more than 0.1% in the fraction of the proteome occupied between young and old worms. Focusing on the 15 proteins that increased the most with age in both datasets, we identified five proteins (VIT-2, VIT-5, VIT-6, PERM-2, TTR-51) that were common between them. All five top proteins contain a signal peptide and are predicted to be secreted extracellular proteins. Notably, three of these proteins (VIT-2, VIT-5, and VIT-6) belonged to the vitellogenin family (Figure 4A, 4B, and 4C) with another member VIT-4 being in the top 50 most upregulated proteins. Therefore, we chose to focus on vitellogenins as candidates for protein buildup components. In addition to the vitellogenins, TTR-51 belongs to a class of transthyretin proteins, a group where five members–TTR-2, TTR-15, TTR-16, TTR-45, TTR-51–were among the top 20 most upregulated proteins with age (Figure S3). Previous work has demonstrated that several members of the TTR family, including TTR-51, are enriched in the secretome of aged animals^44^. Furthermore, human homolog of TTR causes abnormal protein deposits in human tissues and leads to a disease called transthyretin amyloidosis.

**Figure 4.**
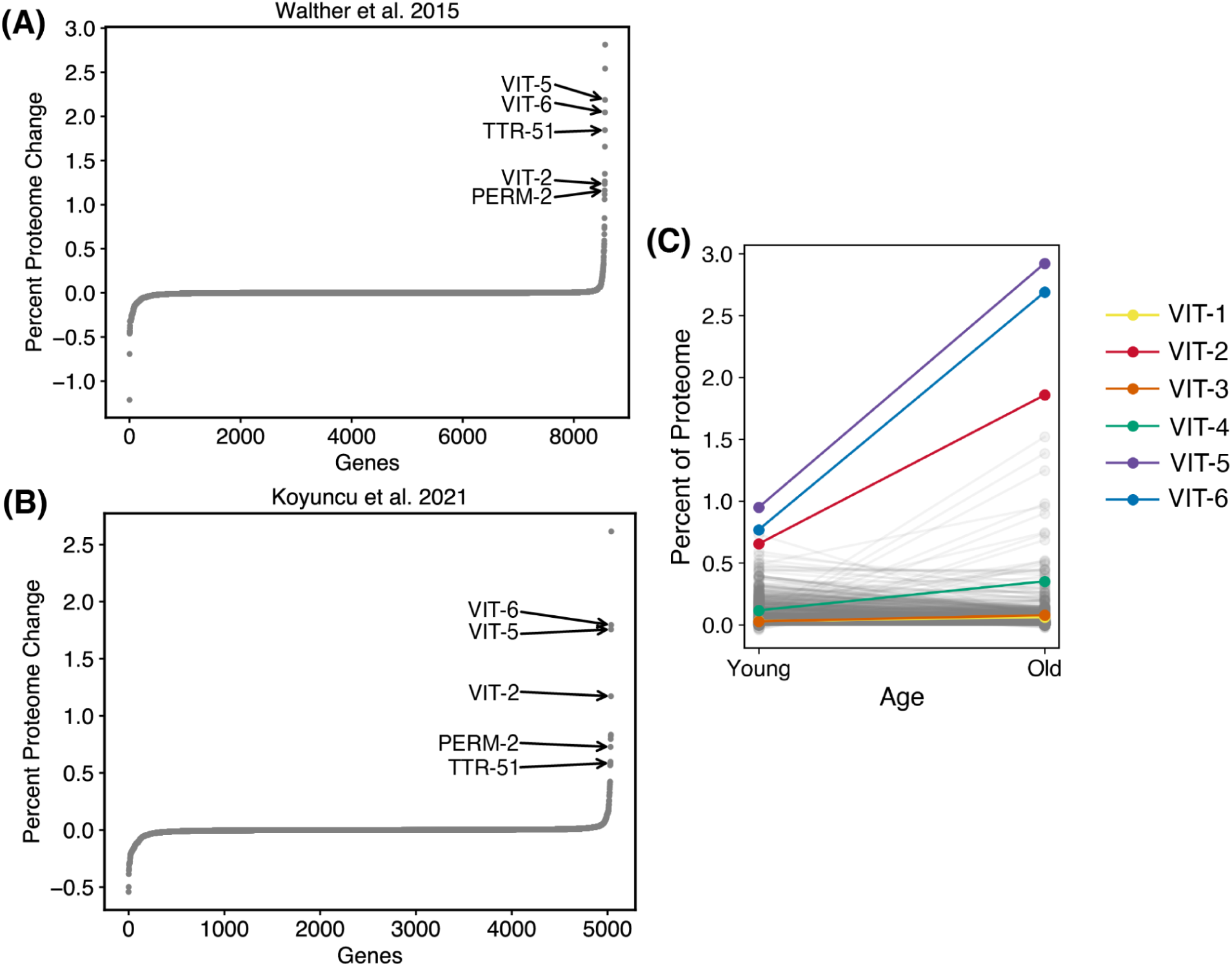
Proteome wide analysis reveals that vitellogenins show the largest age-related increase in absolute abundance **(A)** Ranked distribution of all proteins in Walther et al., 2015 proteomic dataset showing majority of proteins do not change in their percent of proteome occupied with age. Arrows highlight the common 5 genes among the top 15 highest ranked proteins when compared with Koyuncu et al., 2015 proteomics dataset. **(B)** same as (A) but for Koyuncu et al., 2021 proteomics dataset. VIT-2, 5, and 6 are the three most abundant proteins as a fraction of the total proteome with age. Values shown are the average of two independent proteomics datasets.

### Vitellogenins colocalize with the SRS protein buildup and their level is decreased by CR

To determine whether the protein buildup observed in SRS imaging contains vitellogenin proteins, we used strains with GFP and mCherry fused to different vitellogenin proteins. We used three strains containing *vit-2::GFP* ; *vit-6::mCherry*, *vit-2::GFP* ; *vit-3::mCherry*, and *vit-2::GFP* ; *vit-1::mCherry* and performed SRS microscopy with simultaneous 2PF microscopy to examine if the SRS protein buildup signal colocalized with the fluorescent vitellogenin protein signals.

We imaged the strains at two time points: day 1 of adulthood, to confirm the canonical localization of vitellogenin proteins, and day 6 of adulthood, when protein buildup was evident in these older worms. Since *C. elegans* are known to exhibit GFP autofluorescence, we first optimized the photomultiplier tube (PMT) gains for a dynamic range where the low GFP signal detection in N2 worms was balanced against the strong signal from the fluorescent strain worms. These same PMT settings are then used for all 2PF microscopy images (Figure S4 A-D, methods). At day 1 of adulthood, we observed vitellogenin proteins were primarily localized within the intestine, pseudocoelom, oocytes, and embryos, consistent with previous reports of their expected localization in young adults^7^ (Figure 5A, 5C, 5E, and 5G). By day 6 of adulthood, SRS imaging revealed significant protein buildup in pseudocoelomic space, which colocalized with 2PF microscopy signal of vitellogenin proteins—particularly VIT-2 and VIT-6 (Figure 5B, 5D, 5F, and 5H).

**Figure 5.**
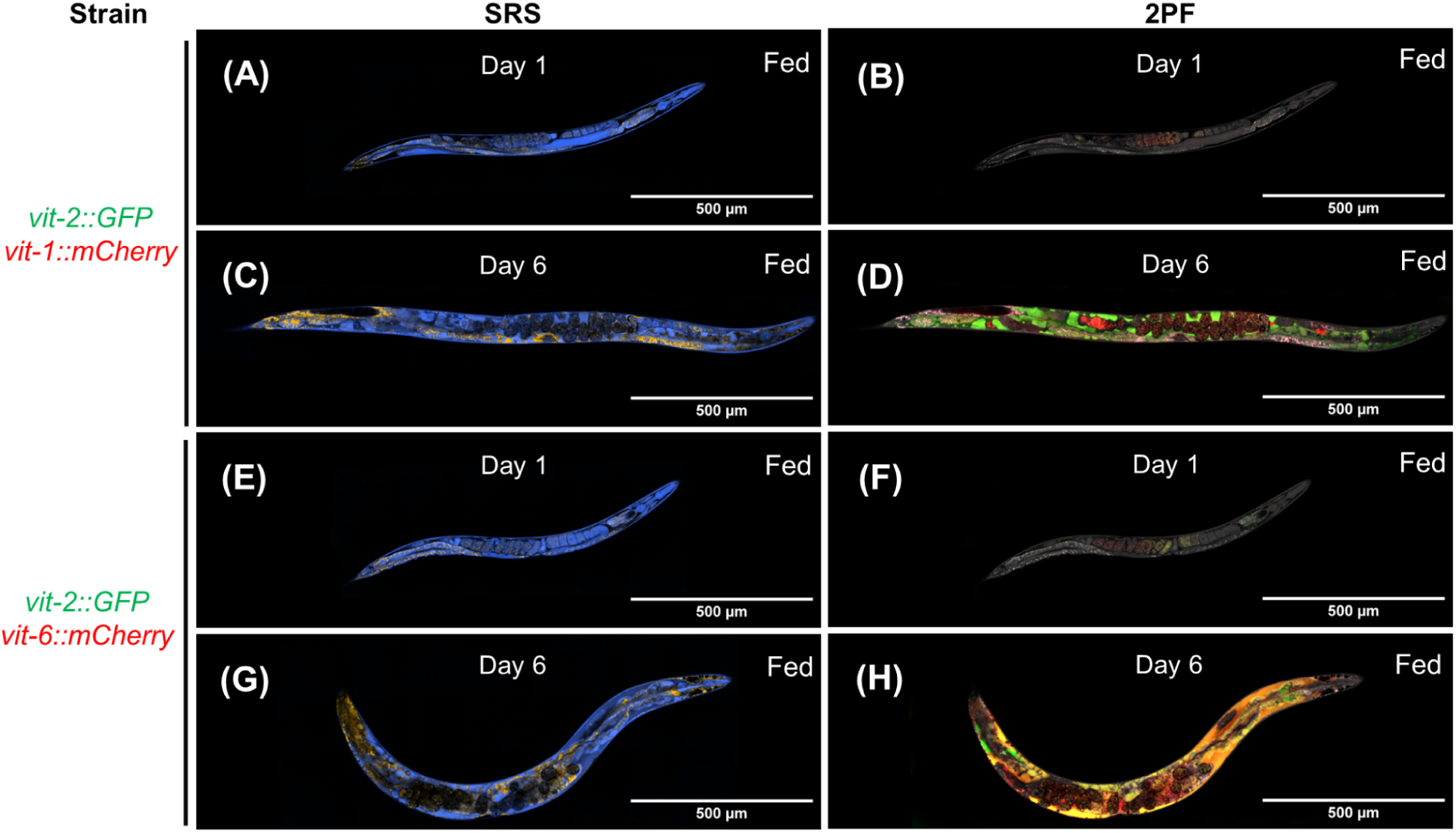
Vitellogenins colocalize with SRS protein buildup **(A)** SRS image of Day 1 adult tagged with vit-1::mCherry and vit-2::GFP **(B)** Composite SRS and GFP and mCherry 2PF image of Day 1 adult tagged with *vit-1::mCherry* and *vit-2::GFP*. SRS CH3 channel shown in gray, GFP in green, mCherry in red. Subsequent composite images are shown in the same color scheme **(C)** SRS image of Day 6 adult tagged with *vit-1::mCherry* and *vit-2::GFP* **(D)** Composite SRS and GFP and mCherry 2PF image of Day 6 adult tagged with *vit-1::mCherry* and *vit-2::GFP* **(E)** SRS image of Day 1 adult tagged with *vit-2::GFP* and *vit-6::mCherry* **(F)** Composite SRS and GFP and mCherry 2PF image of Day 1 adult tagged with *vit-2::GFP* and *vit-2::mCherry* **(G)** SRS image of Day 6 adult tagged with *vit-2::GFP* and *vit-6::mCherry* **(H)** Composite SRS and GFP and mCherry 2PF image of Day 6 adult tagged with *vit-2::GFP* and *vit-2::mCherry*

If vitellogenins are a major component of CR-sensitive SRS protein buildup then CR should reduce vitellogenin levels. To test this, we used strains containing VIT-2 and VIT-6 fused to GFP and mCherry respectively, subjected them to *ad libitum* fed and CR conditions, and measured the expression of these proteins with fluorescence microscopy. Indeed, we found that CR significantly reduced the expression of VIT-2 and VIT-6 (Figure S2). Consistent with this model, previous studies have shown that vitellogenin levels are reduced in *daf-2* mutants^45^.

### Reducing vitellogenesis reduces SRS protein buildup

If the SRS protein buildup consists of vitellogenins, then reducing vitellogenesis should reduce the SRS protein buildup. CEH-60 is a key transcriptional activator of vitellogenesis^13,46–48^. In animals carrying a nonsense mutation in *ceh-60*, the six most downregulated genes in whole-body proteomics are all six vitellogenin proteins (VIT1-VIT6)^46^ . Furthermore, *ceh-60* knockout mutants exhibit a 1,000- to 10,000-fold reduction in vitellogenin gene expression^13^. Therefore, to test whether reduced vitellogenesis leads to decreased SRS protein buildup, we used a strain containing a loss of function mutation in *ceh-60* and performed SRS imaging. Consistent with our hypothesis, we observed that *ceh-60* mutant strains showed reduced protein buildup by day 6 of adulthood (Figure 6B and 6D). At day 12 of adulthood, the total protein levels in fed *ceh-60* mutants were similar to those of CR wild-type animals (Figure 6F, methods).

**Figure 6.**
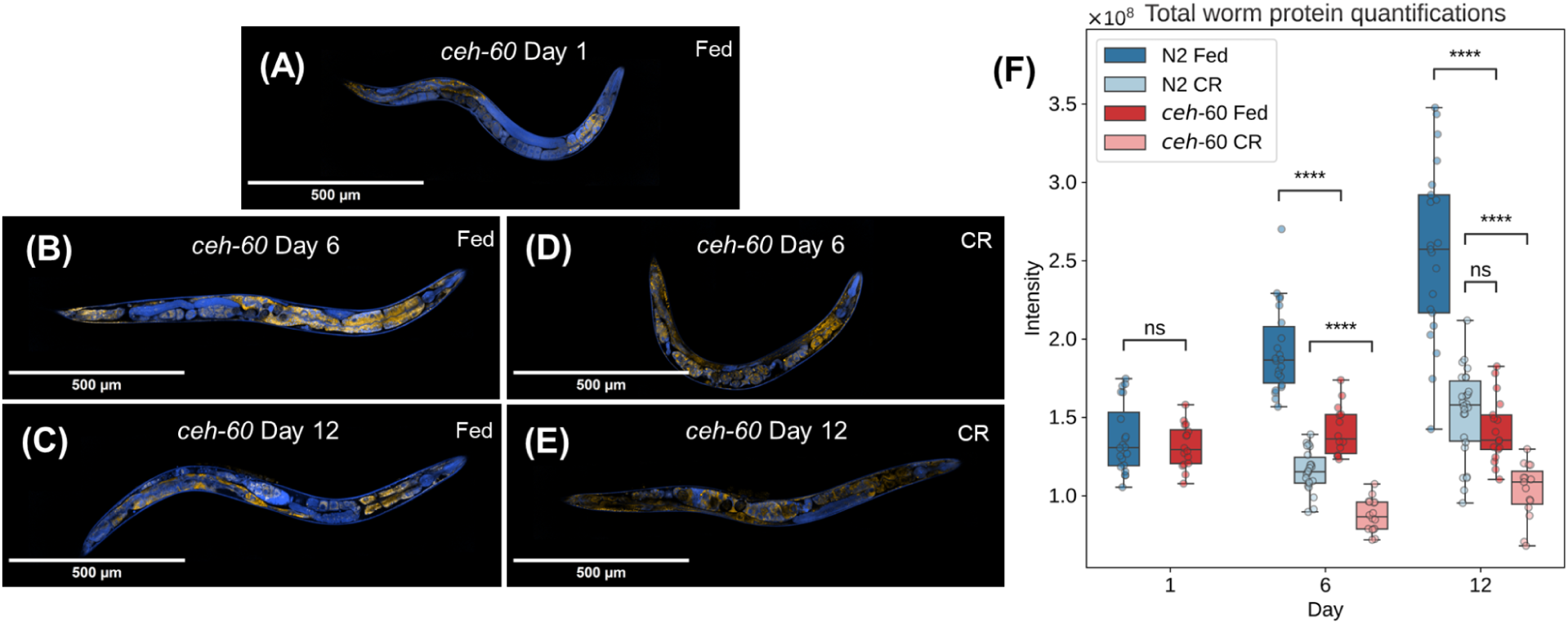
Reducing vitellogenesis reduces SRS protein buildup **(A)** SRS image of Day 1 Adult *ceh-60* KO, fed **(B)** SRS image of Day 6 Adult *ceh-60* KO, fed **(C)** SRS image of Day 12 Adult *ceh-60* KO, fed **(D)** SRS image of Day 6 Adult *ceh-60* KO, fed **(E)** SRS image of Day 12 Adult *ceh-60* KO, CR **(F)** Quantification of total protein of N2 and *ceh-60* KO fed and CR across Day 1, 6, and 12 *, **, ***, and **** indicate p < 0.05, 0.01, 0.001, and 0.0001 (Welch’s t-test), respectively. Horizontal bars of the boxes represent maximum, 0.75, 0.5, 0.25, minimum, respectively. Outliers are determined by whether the point deviates 1.5 times the interquartile range (0.75 quartile - 0.25 quartile), from either quartile.

### Reducing vitellogenesis extends lifespan of fed but not CR worms

Since CR reduces vitellogenin buildup and decreasing vitellogenesis by itself is known to extend lifespan we asked if CR extends lifespan by decreasing vitellogenesis^9,11–13,37^. To test this, we conducted a lifespan epistasis experiment using the *ceh-60* loss of function mutant, previously shown to exhibit lifespan extension under fed conditions^13^. While loss of *ceh-60* led to a robust >60% lifespan extension under *ad libitum* fed conditions, there was no further extension of lifespan under CR compared to that of wild-type N2 worms (Figure 7A). Similarly, we found that loss of *ceh-60* provides protection under heat stress conditions (30°C) like CR – and combining CR and loss of *ceh-60* together provided no additional protection (Figure S6). To further confirm that reducing vitellogenesis is working non-additively with CR to extend lifespan, we used a strain containing loss of function mutations in all six vitellogenin genes. Consistent with *ceh-60* findings, we found that under *ad libitum* fed conditions, loss of all six vitellogenin genes led to >60% lifespan extension and combining loss of vitellogenins with CR provided no further extension of lifespan compared to wildtype CR (Figure 7B). CR still further extended the lifespan of both loss of function *ceh-60* and *vit-1-6(null)* worms, suggesting that CR may be extending lifespan through additional mechanisms besides reducing vitellogenesis.

**Figure 7.**
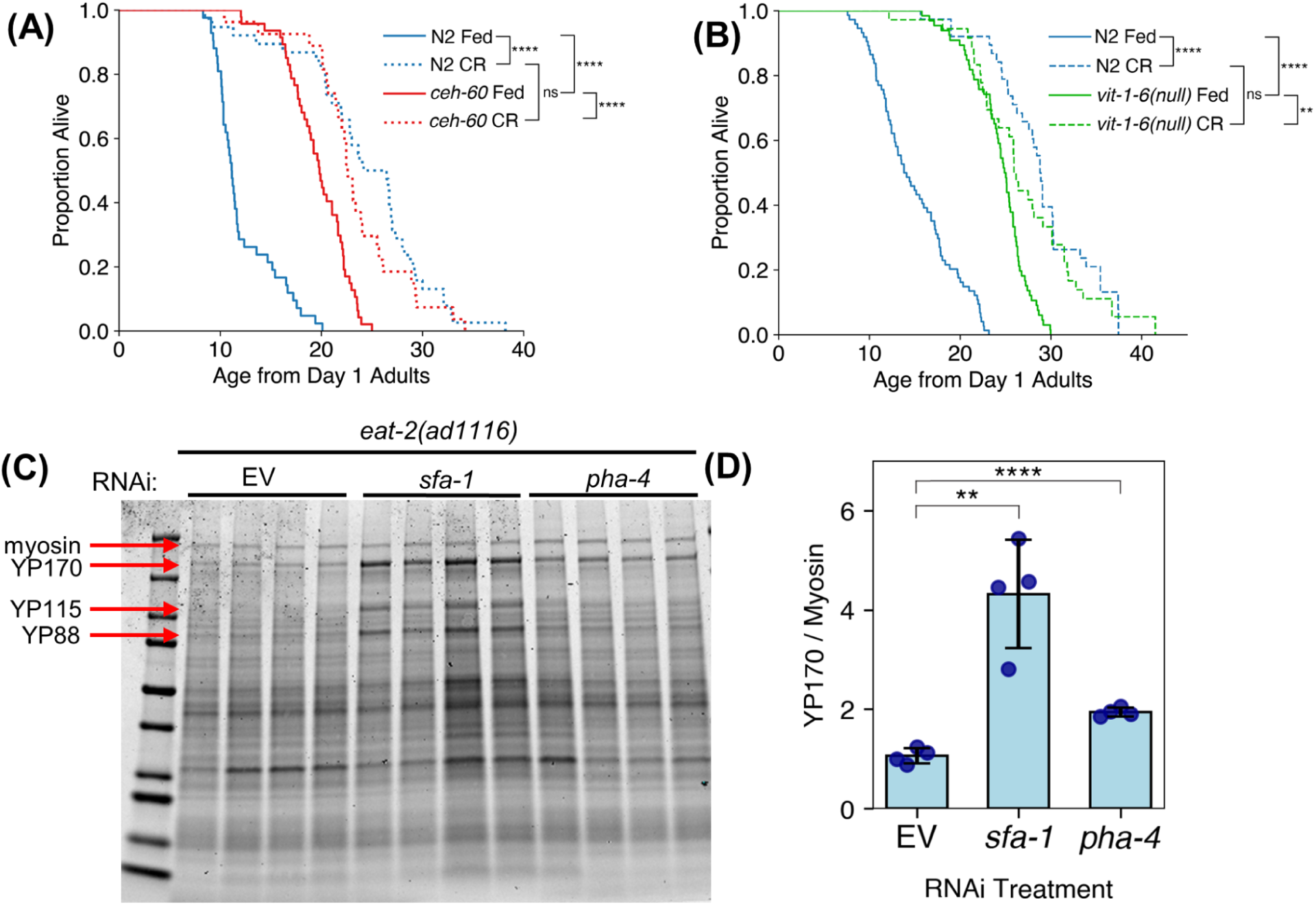
Reducing vitellogenesis extends lifespan of fed but not CR worms **(A)** Survival curves of N2 and *ceh-60* KO under both fed and CR conditions **(B)** Survival curves of N2 and *vit-1-6(null)* under both fed and CR conditions **(C)** Yolk protein measurement for *eat-2(ad1116)* worms with RNAi for genetic regulators of CR. EV = Empty Vector and represents control Quantification of YP170 levels normalized to myosin for (C). Data is plotted as mean ± SD. ** indicates p < 0.01 and **** indicates p < 0.0001

### Genetic regulators of CR affect vitellogenesis

Genetic factors essential for CR-mediated lifespan extension have been identified, including *sfa-1* and *pha-4*^2,49–52^. Therefore, we asked whether vitellogenins act downstream of these genetic factors. If true, knocking down (KD) these genes during CR – thereby preventing CR-mediated lifespan extension – should result in elevated vitellogenin levels compared to control CR worms. Observing this would provide independent evidence supporting the model that CR and reduced vitellogenesis are working in the same pathway. To test this, we used feeding RNAi to KD *sfa-1* and *pha-4* in CR conditions^49–51^. We found that RNAi KD of these factors led to higher vitellogenin levels during CR compared to their corresponding CR control, suggesting that vitellogenins may lie downstream of these genes to regulate the longevity response to CR (Figure 7C and 7D). Taken together, these findings support a model in which CR extends lifespan by downregulating vitellogenesis.

## DISCUSSION

### Stimulated Raman scattering microscopy is a powerful tool to study aging

Aging is associated with a loss in protein homeostasis but how proteins change both in abundance and spatial location across an animal remains less well understood ^1,53–56^. To characterize these changes, we used SRS microscopy to capture changes in protein and lipid levels within the entire body of *C. elegans*, across its lifespan, and with two of the most robust longevity interventions – CR and reduced IIS. Analysis of these images revealed a striking buildup of protein in the pseudocoelomic space of aged *C. elegans*. We sought to identify the composition of these proteins that were building up and determine whether they contributed to the regulation of aging rates. Following analysis of proteomics data from aging worms, we identified the protein build up observed in the SRS microscopy images could be attributed to vitellogenins or yolk proteins.

In addition to visualizing protein build up, the SRS images acquired in this study reveal many interesting features of aging and caloric restriction. Our study provides a comprehensive database containing over 12,000 images of 5166 FOV’s across 355 whole worms, detailing spatial and concentration changes in proteins and lipids with age, dietary interventions, and genetic manipulations which we hope will be a valuable resource to the aging community. Here we have shown extensive SRS imaging of over 350 whole worms at protein and lipid bands, but SRS offers many more capabilities that are useful towards whole animal imaging. For example, the images shown here represent a thin slice in Z through the middle of the worms with ∼1μm resolution due to the multiphoton optical sectioning effect afforded by SRS microscopy. While this is an advantage for observing features in high resolution in the axial direction, 3D images might offer a more complete picture of the worm. 3D SRS images are of course possible, but given the thickness of the worms and necessary grid stitching of images in XY, imaging times can become cumbersome. We do present, however, a few examples of 3D images acquired that demonstrate the expected exoskeleton features such as the cuticles of the worm (Figure S7). Additionally, given the nondestructive imaging of SRS, longer term time-lapse imaging of worms is also feasible with temporal resolution of seconds for a given field of view.. We also note the existence of many interesting chemical and metabolic features observable in other parts of the Raman spectrum. While the lipid and protein peaks presented here allow for observation of the vast majority of anatomical features within the worms, they do not offer high molecular specificity as has been demonstrated in other works that image in the fingerprint of the Raman spectrum^22,26,28^ or use carbon-deuterium pulse-chase labeling to identify metabolic turnover and lipogenesis^31,57^. Hyperspectral imaging is another potential future direction for more advanced analyses of internal molecular content here.

### Vitellogenesis is an important determinant of lifespan in C. elegans

Multiple studies have implicated vitellogenesis as a driver of aging in *C. elegans*, and our study now adds to this growing literature. RNAi of multiple vitellogenin genes or loss of function in *ceh-60*, a positive regulator of vitellogenesis, have been shown to extend lifespan^9–13^. This lifespan extension appears independent of germline loss, as defects in vitellogenesis do not reduce brood size^13,47^ .

SRS microscopy revealed a reduction in protein buildup during CR, prompting us to investigate whether reduced vitellogenesis is working through a shared pathway with CR to extend lifespan. Previous studies suggest this could be the case. Lower mRNA levels of all six vitellogenin genes and reduced levels of the vitellogenin protein complex YP170 in *C. elegans* were observed in a genetic model of CR^12,58^. In addition, overexpressing a single vitellogenin gene, *vit-2*, shortened the lifespan of the same CR model^12^. Consistent with these findings, we used fluorescence microscopy on worms expressing GFP and mCherry fused to vitellogenin proteins and confirmed that CR reduces vitellogenin levels. Using lifespan epistasis experiments, we demonstrated that lifespan extension from reduced vitellogenesis is non-additive with CR, suggesting that these two longevity pathways are working through a similar pro-longevity pathway.

Adult *C. elegans* dedicate significant resources to vitellogenesis, making it a natural candidate for a process contributing to aging^6,7^. It has been estimated that vitellogenin proteins occupy a striking ∼30-40% of the total worm protein by day 5 of adulthood, and their accumulation has been documented using fluorescent tags and immuno-EM^7,9,59–61^. Our study adds to this observation using an unbiased orthogonal label-free method to find highly increased proteins with age. We further utilized a total protein approach based on previously published proteomic measurements to show that vitellogenin proteins are the most abundant class of enriched proteins with age in *C. elegans*^38,43^. We then experimentally confirmed that vitellogenins were localized inside the regions of protein buildup observed by SRS in aged worms.

### Vitellogenesis and the Antagonistic Pleiotropy model of aging

A framework to understand why *C. elegans* continue to expend energy to produce vitellogenins despite its negative effects is the “Antagonistic Pleiotropy” theory^10,62–64^. This theory posits that evolutionary pressures select for genes that provide early-life fitness benefits even if they impose late-life costs. This would occur if the early-life benefits are greater than the late-life costs to the individual. Previous studies have shown that vitellogenins offer early-life benefit by supplying nutrients to developing oocytes and enhancing the survival of L1 larvae during periods of food scarcity^65,66^. However, their continued production later in life could then impose a cost – expending energy for their continued production that leads to protein aggregation and contributes to aging and death.

### Parallels between vitellogenesis in C. elegans and apolipoprotein B in humans

Vitellogenins are conserved in all oviparous animals, from nematodes to birds, and even monotremes within the mammalian lineage^67^. While humans have evolved the use of a placenta to deliver nutritional resources to their offspring, the sequence and functional similarity of vitellogenins to ApoB suggests our finding may be applicable to placental mammals as well^15^. Similar to our observation that CR reduces vitellogenin levels in *C. elegans*, CR in mice has been shown to reduce hepatic ApoB expression^12^. Humans who are under a voluntary CR regimen also show reduced circulating levels of LDL compared to age-matched controls^16^, suggesting a similar modulation of ApoB expression. Furthermore, increased circulating ApoB levels in humans is associated with age and age-related diseases such as cardiovascular disease^14,17,18^.

## METHODS

### Chemicals and Materials

| **Chemicals and Materials** | **Source** | **Identifier** |
| --- | --- | --- |
| Cholesterol | Sigma | C8667 |
| LB Broth, Miller | Fisher Scientific | BP1426-500 |
| KH<sub>2</sub>PO<sub>4</sub> | Sigma | P9791 |
| K<sub>2</sub>HPO<sub>4</sub> | Sigma | 7758-11-4 |
| MgSO<sub>4</sub> | Strem | 93-1255 |
| Bacto-Peptone | Fisher Scientific | 211677 |
| Bacto Agar | Fisher Scientific | 214030 |
| NaCl | Sigma | 567440 |
| Palmitic Acid | Sigma | P0500 |
| FUDR | RPI | F10705 |
| Kanamycin | Gold Bio | K-120-5 |
| Carbenicillin | Gold Bio | C-103-5 |
| Tetracycline Hydrochloride | Sigma | T7660 |
| Sodium Azide | Sigma | S2002 |
| Agarose | BioRad | 1613101 |
| IPTG | Gold Bio | I2481C |
| Imperial Protein Stain | Fisher Scientific | 24615 |
| Lauryl sulfate (SDS) sodium salt | Sigma | L-3771 |
| Dulbecco's PBS | Sigma | D8537 |
| 4x Laemmli Sample Buffer | BioRad | 1610747 |
| 2-Mercaptoethanol | Sigma | M3148 |
| 4–20% Mini-PROTEAN® TGX™<br>Precast Protein Gels, 15-well, 15 µl | BioRad | 4561096 |
| Tris-Glycine-SDS Running Buffer<br>(10X) | Boston BioProducts | BP-150 |
| Slide microscope single plain<br>75x25x1mm | Thomas Scientific | 1227Z74 |
| Cover glass slips, 22x22mm #1,<br>microscope slide | VWR | 48366-067 |
| 50 x 9mm Petri Dish Plates | Fisher Scientific | 8757105 |
| Slide microscope single plain<br>75x25x1mm | VWR | 16004-422 |
| Dimethyl Sulfoxide | Fisher Scientific | BP231-100 |
| 1x PBS | Corning | 21040CV |
| Type N Immersion Liquid | Leica | 11513860 |

#### Worm strains and culture

The following C. elegans strains were obtained from the Caenorhabditis Genetic Center, funded by the NIH Office of Research Infrastructure Programs (P40 OD010440): N2(Bristol) wildtype, MQD2774 (vit-6(hq486[vit-6::mCherry]) IV; vit-2(crg9070[vit-2::gfp]) X), MQD2775 (vit-3(hq485[vit-3::mCherry]) vit-2(crg9070[vit-2::gfp]) X), MQD2798 (vit-2(crg9070[vit-2::gfp]) vit-1(hq503[vit-1::mCherry]) X), and VC988 (ceh-60(ok1485) X). The CB1370 (daf-2(e1370) III) strain was a gift from A. Dillin. DLS1004 (vit-6(rhd332[S320F, W322*]) IV; vit-5(rhd322[T391E, L392F, A393*]) vit-4(rhd321[T391E, L392F, A393*]) vit-3(rhd320[T391E, L392F, A393*]) vit-2(ok3211) vit-1(ok2616) X) was a gift from R. Dowen.

Worms were routinely grown at 20°C with the exception of CB1370 which was grown at 15°C to prevent dauer formation till they reached L4. Worms were grown on 60mm plates with standard nematode growth media (NGM) containing 1 mM CaCl2, 5 µg/mL cholesterol, 25 mM KPO4 pH 6.0, 1 mM MgSO4, 2% (w/v) agar, 0.25% (w/v) Bacto-Peptone, 51.3 mM NaCl and seeded with E. coli (OP50). E.coli bacteria was inoculated overnight in Luria Broth (LB) Media and grown at 37°C, after which 250ul of the culture was seeded onto plates and left to dry at least overnight before worms were transferred onto the plates.

#### Lifespan Machine Plate preparation

Separate plates were made to be compatible for measurement with the “Lifespan Machine” which worms are transferred to on day 6 of adulthood. Nematode Growth Media was made by first autoclaving 0.25% (w/v) Bacto-Peptone, 51.3 mM NaCl, and 1.7% (w/v) agar media. After cooling to 60C the following components were added to a final concentration of 5µg/mL cholesterol, 25 mM KPO4 pH 6.0, 1 mM MgSO4, and 40uM of FUDR. 8mL of the following media was poured into 50x9mm Falcon (Fisher Scientific Cat#8757105) plates in a Tissue Culture hood and left for 45 minutes to solidify. 200µl of 10mg/mL Palmitic Acid dissolved in EtOH was then added as a ring by pipetting 100µl and swirling quickly, followed by another 100µl immediately on top to create a thicker ring layer around the outside of the plate and left to dry for an additional 2 hours with no lids. The palmitic acid ring was used to prevent caloric restricted worms from escaping. Plates were then capped tightly and stored in 4°C until one day before worms were transferred to them.

#### Lifespan Assays

All lifespan experiments were conducted at 20°C, with the exception of CB1370 and the corresponding N2 control which were grown at 15°C from egg till L4, and then grown at 20°C for the rest of their life. All worms were synchronized through a timed egg lay. Starting from day 1 till day 6 of adulthood, all worms were transferred daily to fresh plates that were always made 1 day before worms were transferred onto the plate. 7 worms were transferred per plate to prevent running out of food for CR conditions.. To make these plates, 60mm plates were made with standard nematode growth media as described under the “Worm strains and culture” protocol and were seeded with either 250µl of OD600 of 20 for “*ad libitum* fed” conditions or OD600 of 0.1 for “CR” conditions. Following seeding, plates were incubated at 37°C for 1 hour, then spiked with an antibiotic and drug cocktail for a final concentration of 50uM FUDR5-fluoro-2′-deoxyuridine (FUDR), 50µg/mL kanamycin, and 100µg/mL carbenicillin following the sDR protocol to prevent the hatching of eggs and the growth of bacteria^68^.

At day 5 of adulthood or one day prior to transferring worms to the “Lifespan Machine”, Lifespan Machine Plates were seeded with either 200µl of OD600 of 20 or 200µl of OD600 of 0.1 for “ad libitum” or “CR” conditions respectively. Seeded plates were incubated at 37°C for 1 hour and then spiked to a final concentration of 50µg/mL Kanamycin and 100 µg/uL carbenicillin per plate to prevent the growth of bacteria. At day 6 of adulthood, worms were transferred to these seeded Lifespan Machine Plates and loaded into the “Lifespan Machine”. The machine is set to scan once every 15 minutes, so that every plate gets imaged once per hour for the duration of their life. Using a trained worm detection model and movement analysis their survival was recorded and analyzed as previously described^69^.

#### Thermotolerance Assay

Worms were synchronized via egg lay and grown till day 1 adults at 20°C. At day 1 of adulthood, worms were transferred to either fed or caloric restricted plates described in the “Solid Dietary Restriction” protocol. Worms were transferred daily to freshly made fed or caloric restricted plates from day 1 till day 4 of adulthood. At day 4 of adulthood, worms were placed into an elevated 30°C temperature for the rest of their life and survival was scored with a fully automated lifespan machine.

#### Solid Dietary Restriction Protocol

Caloric Restriction assays were performed as previously described with some modifications^68^. Worms were synchronized via egg lay and grown till day 1 of adulthood. At day 1 of adulthood, worms were transferred to fed or caloric-restricted plates. Plates were prepared by culturing E. coli OP50 and concentrated and the absorbance at OD600 was measured. The bacterial stock was then diluted to either an OD600 of 20 for fed plates or 0.1 for caloric-restriction plates. 250µl of OD20 or OD0.1 was seeded onto 6cm standard NGM plates and incubated at 37°C for 1 hour. Plates were then spiked with a final concentration per plate of 50µM 5-fluoro-2′-deoxyuridine (FUDR), 50µg/mL kanamycin, and 100µg/mL carbenicillin. Plates were made one day prior to transferring worms to the plates. Worms were transferred daily to new plates from day 1 till day 6 of adulthood.

#### C. elegans sample preparation for SRS imaging

75x25x1mm microscope slides (Thomas Scientific Cat# 1227Z74) for imaging were prepared by making a freshly flattened 2% agarose pad for every round of imaging. Animals were first picked away from any food before being immobilized in 8ul of 100mM Sodium Azide dissolved in M9 then sealed beneath a coverglass (VWR Cat# 48366-067). 6 -10 animals were immobilized per slide and were immediately imaged. Fresh batches of agarose pads and worms were immobilized and prepared every 2 hours to ensure the samples that were imaged were as fresh as possible till a sufficient sample size was acquired per condition.

#### SRS and 2PF microscopy

SRS microscopy coherently excites chemically-specific Raman vibrational modes of molecules based on a multiphoton interaction of two lasers at the focus of an objective. The wavelengths of the interacting lasers are chosen such that the difference in photon energies between the two beams is “resonant” (i.e. equal to) the energy of a particular Raman vibration of interest. These lasers are then point scanned through the back of the objective to create an image of the microscopic field of view (FOV) where each pixel in the image corresponds in value to the strength of SRS signal. Under the assumption of fixed laser powers at focus, the strength of SRS signal (and thus pixel values) is linearly related to the concentration of molecules within the focal volume at that vibrationally resonant mode. Thus, SRS microscopy images can be thought of as “concentration maps” of Raman active molecules for a given vibrational mode within a field of view. By changing interacting wavelengths of the lasers, different Raman vibrational transitions (i.e. different molecules or molecular moieties) can be interrogated. A more thorough description of the SRS microscopy and its quantitative implications can be found in Potma^70^ and in Manifold and Fu^71^.

Our set up utilizes a broadband spectral focusing arrangement with high density glass rods to impart matched chirp on the pulses of the two beams as described previously^72–74^. Here, laser wavelengths of 799 nm and 1040 nm from an Insight DS+ are chosen for the pump and Stokes beams, respectively. The two imaged Raman transitions (2850 cm^-1^ and 2920 cm^-1^) are then tuned between by changing the temporal delay between the pulses via a computer controlled retroreflector stage (Newport FCL 200) that has been calibrated in the spectral region afforded by the broadband pulses (∼2800-3050 cm^-1^). The laser beam powers at focus were 20 mW and 37 mW for pump and Stokes respectively. The beams are raster scanned through an Olympus Fluoview IX83 Microscope equipped with a UPlanSApo 60x 1.2 NA water immersion objective and Nikon 1.4 NA oil immersion condenser. This affords our images an ultimate lateral spatial resolution of ∼400 nm. The transmitted light is then filtered of the 10.28 MHz modulated Stokes beam and the pump is detected on a 64V-biased silicon photodiode (Hamamatsu S3590-08). The output of the photodiode is then sent through a bandpass filter (mini-circuits) and voltage amplifier (Electro Optical Components DHPVA-201) to a lock-in amplifier (Zurich Instruments HF2LI) triggered to the same 10.28 MHz modulation driving frequency to detect the imparted SRS signal. The phase and lock-in gain of the SRS signal is set to be consistent across all experiments for viable comparison of relative signal strengths. The output voltage of the lock-in is then sent to the Olympus analog-to-digital converter to integrate the SRS signal to the FluoView software where images are acquired. Python scripts are used to control the movement of the delay stage, XY stage, and acquisition of images.

For the imaging of worms with fluorescent reporters present, the Olympus microscope’s equipped non-descanning multiphoton photomultiplier tubes (PMT) were used to simultaneously capture the 2PF images. A 570 nm long-pass dichroic separates the GFP and mCherry fluorescence in the back direction from the objective, then they are respectively filtered by 520/30 nm and 640/50 nm bandpass filters before detection on the PMT’s. PMT gains were calibrated based on the strongest observed fluorescence in worms with an upper buffer to maximize the dynamic range. Once gains were set, they were held consistent across all imaged worms to allow for relative signal comparison.

For the acquisition of worm images, the focus is set to the middle of the worm as observed by the junction between the pharynx and intestinal tract. Then the minimum circumscribing rectangle is calculated around a worm in XY stage coordinates. A python script then creates a grid map of XY stage positions at which to acquire SRS images (protein + lipid), then initiates image taking through the grid sequentially. Images were acquired at 512 x 512 pixel resolution with a 10 μs pixel dwell time (∼2.6 seconds per image). Whole live worms were imaged in 2 - 9 minutes depending on the size and orientation of the worm.

#### Fluorescence stereomicroscope imaging

MQD2774 (vit-6(hq486[vit-6::mCherry]) IV; vit-2(crg9070[vit-2::gfp]) X) worms were prepared following the “Solid Dietary Restriction” protocol and aged till day 5 of adulthood. On day 5, animals were removed from the bacterial lawn and anesthetized in 100mM sodium azide and lined up on an unseeded 60mm NGM plate. Animals were imaged using an M250FAsteroscope (Leica) under the respective fluorescence.

#### Image field-correction and stitching

In each experiment, prior to imaging of worms, an image of 25% DMSO solution is taken. This image is flattened with a gaussian convolution (blur). The inverse of the image is then multiplied by the original maximum, creating the field-correction multiplier image. Every worm image from the same experiment was multiplied by this correction image, to compensate for the artificial intensity difference between center and edge.

Acquired 4D image hyperstacks (X,Y,C,T) are rearranged and separated into labeled individual 3D tiles (X,Y,C). With the grid information (X and Y count) recorded during the microscope experiments, the worms are stitched using the ImageJ grid stitching plugin.^75^ Composite SRS or fluorescent microscopy images are generated and rescaled to ease the check of the stitching result.

#### Image segmentation

Each worm is segmented in a hybrid approach.^76^ Firstly, the image is coarsely segmented using a custom Python script. Before segmentation, there is a morphologically reconstruction step using the erosion method. The seed image is identical to the original image, except the edge rectangle assigned with the maximum value from the original image. After the reconstruction, a hard otsu thresholding is applied. Small objects are removed, followed by a closing operation, to make sure the worm body mask has no holes or other artifacts.

Manual adjustment of the masks are performed to further improve the accuracy of downstream analyses. For instance, worms carrying embryos introduce additional regions to the segmentation mask. Every mask is overlaid on the original image with a transparency, and manually corrected. The correcting operations include removing the eggs, and fixing the missing area due to low signal and hard thresholding.

#### Worm morphology characterization

The worm morphology needs to be characterized before quantitative analysis. This means to identify the spline of the worm, as well as its head and tail direction. To achieve this efficiently, a custom analysis procedure is performed as follows.

The border of the worm is found by selecting the largest contour line.^77^ This line is then converted to an ordered path. To obtain head and tail endpoint locations on the path, interspaced points on the path are selected, to calculate the angle between each adjacent line segment pair. By choosing an appropriate sampling rate, the two endpoints on the paths can be identified.

The border path of the worm is then cut into halves by the two endpoints. The two halves are resampled with interpolations, and averaged together into the spline of the worm. A fixed quantile (0.1) point of the spline is used to divide the original mask into two halves. This divided mask is then overlaid on the original image, and manually checked if it is closer to the head. After the checking, the two endpoints are classified into head and tail endpoints. The head part of the divided mask is then saved as a separate anterior worm mask.

#### Quantification and statistics

With this annotated data, including the morphological characterization, quantifications can be performed as follows.

Each worm image has two channels, corresponding to the two Raman transitions stated in section “SRS and 2PF microscopy”. As described in the results, the 2850 cm^-1^ channel then serves as the proxy for lipid content, thus named “lipid channel”. The lipid channel is normalized to the 2920 cm^-1^ channel image, to let the two maximal intensity values match. After the normalization, the lipid channel is subtracted from the 2920 cm^-1^ image, and the new image is called “protein channel”. This is because not only protein but also lipid contributes to the CH3 peak (2920 cm^-1^).

The 2-channel sum intensity values of worms are then calculated, as well as the area of the masks. Derivatives of the sum and the areas include average intensities of protein/lipid channels, ectopic area, and ectopic lipid sum. These intensity values are then combined with the metadata including worm age, genotype, caloric restriction condition, and experimental parameters.

For boxplots, bars in each box show the maximum, minimum, median, 25% and 75% percentile of the distribution. Outliers are determined by whether the point deviates from 1.5 times the interquartile range.

Welch’s t-test was performed on multiple genotypes or genotype & condition pairs. For lifespan experiments, survival curves were analyzed using the Log-rank test. Thresholds for p-values were set as follows: 0.05 for significant (one star), 0.01, 0.001 and 0.0001 for two, three and four stars, respectively.

#### Feeding RNA interference

All RNAi constructs came from the Ahringer library and their plasmids were purified and sequenced to verify matching gene targets before use. RNAi experiments were carried out using NGM plates supplemented with 100µg/mL carbenicillin and 1mM IPTG (termed “RNAi NGM plates”). RNAi bacteria were cultured overnight at 37°C in 30mL of LB containing 100µg/mL Ampicillin and 10µg/mL tetracycline. The RNAi bacteria were centrifuged and supernatant was removed and replaced with 10mL of M9. The bacterial concentration of all RNAi strains was measured using absorbance at 600nm and equalized to 2.5 OD600 to control for changes in food density. 250ul of each RNAi strain was seeded per “RNAi NGM plate” and left for at least 48 hours before use at room temperature. *sfa-1* RNAi was started from egg lay, while *pha-4* was started from day 1 of adulthood.

Starting from day 1 of adulthood, worms were transferred to RNAi NGM plates with FUDR to prevent progeny hatching. To make these plates, RNAi NGM plates were seeded with RNAi bacteria with matching OD600 values for at least 48 hours before being spiked with 100µl of FUDR directly to the center of the bacterial lawn to make a final concentration of 50uM of FUDR per plate. Plates were left on the bench to dry for at least 24 hours before worms were transferred onto the plate. Since FUDR can affect the effectiveness of RNAi, we spiked an additional 100µl of IPTG onto the center of the bacterial lawn on the same day we planned to transfer worms to the plate, (for a final concentration of 2mM IPTG per plate) and let the plate dry for an additional 2 hours before transferring the worms to the plate.

#### Yolk Protein Measurements

Yolk Proteins were quantified by running worm lysates on a protein gel and staining with an Imperial Protein Stain for proteins (Thermo Cat#24615). Lysis buffer was prepared by making a master mix containing 375ul of 5% SDS in PBS with 125µl of 4x Laemmli Buffer with 2-mercaptoethanol, and aliquoting 20µl of this mixture. 20 day 4 adult worms were picked into 20µl of lysis buffer and immediately snap frozen in liquid nitrogen. They were then boiled at 99C for 10 min in a thermocycler, then centrifuged, and boiled again for another 10 min. This was repeated such that tubes were boiled for a total of 30 min at 99°C. 5ul of this lysis was then loaded onto a 4-20% Mini-PROTEAN® TGX™ Precast Protein Gels, 15-well. The gel was run at 100V for 20 min followed by 200V for 35 minutes. The gel was washed for 15 minutes in milliQ water (replaced with fresh milliQ water every 5 minutes), and then stained with the Imperial Protein stain for 1 hour. The gel was then washed overnight in milliQ water before imaging. Gels were imaged using a Li-COR imager (LI-COR Biosciences). Bands were quantified using a densitometric analysis with ImageJ.

#### Proteomic data analysis

We utilized a total protein approach and previously published measurements of the absolute quantification of proteins to estimate the fraction of the proteome occupied by each protein detected in the proteomics dataset over age^38,43^. We only included studies that utilized a label-free DIA (Data-Independent Acquisition) approach. The total protein approach assumes that a protein’s abundance within a cell’s proteome is proportional to that protein’s mass spectrometry signal (MS) over the total MS signal.

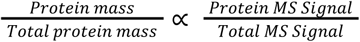

To reduce noise from the proteomics data, we first calculated the best fit line of the fractional occupancy of each protein identified in the proteomics dataset over age. From the best fit calculation, we subtracted the fraction the protein occupied at the oldest age minus the fraction occupied at the youngest age (Day 1 of adulthood) (Figure 4A). This gave us a quantifiable value for each protein as a percent change in the proteome occupancy with age. For each proteomics dataset, we then ranked all proteins from highest to lowest change in proteome occupancy. Taking the top 15 highest ranked proteins found in each dataset, we then merged the two to find the common proteins (Figure 4C and 4D).

## Acknowledgements

Research reported in this publication was supported by the National Science Foundation under award number 1845623 (A.S.), the National Institutes of Health under award number R35GM124916 (A.S.) and DP2GM132933 (D.V.T), and the Harvey and Leslie Wagner Foundation. A.S. is a Chan Zuckerberg Biohub Investigator. We thank Robert Dowen for sharing the *vit* KO strain with us.

## SUPPLEMENT

**Figure S1.**
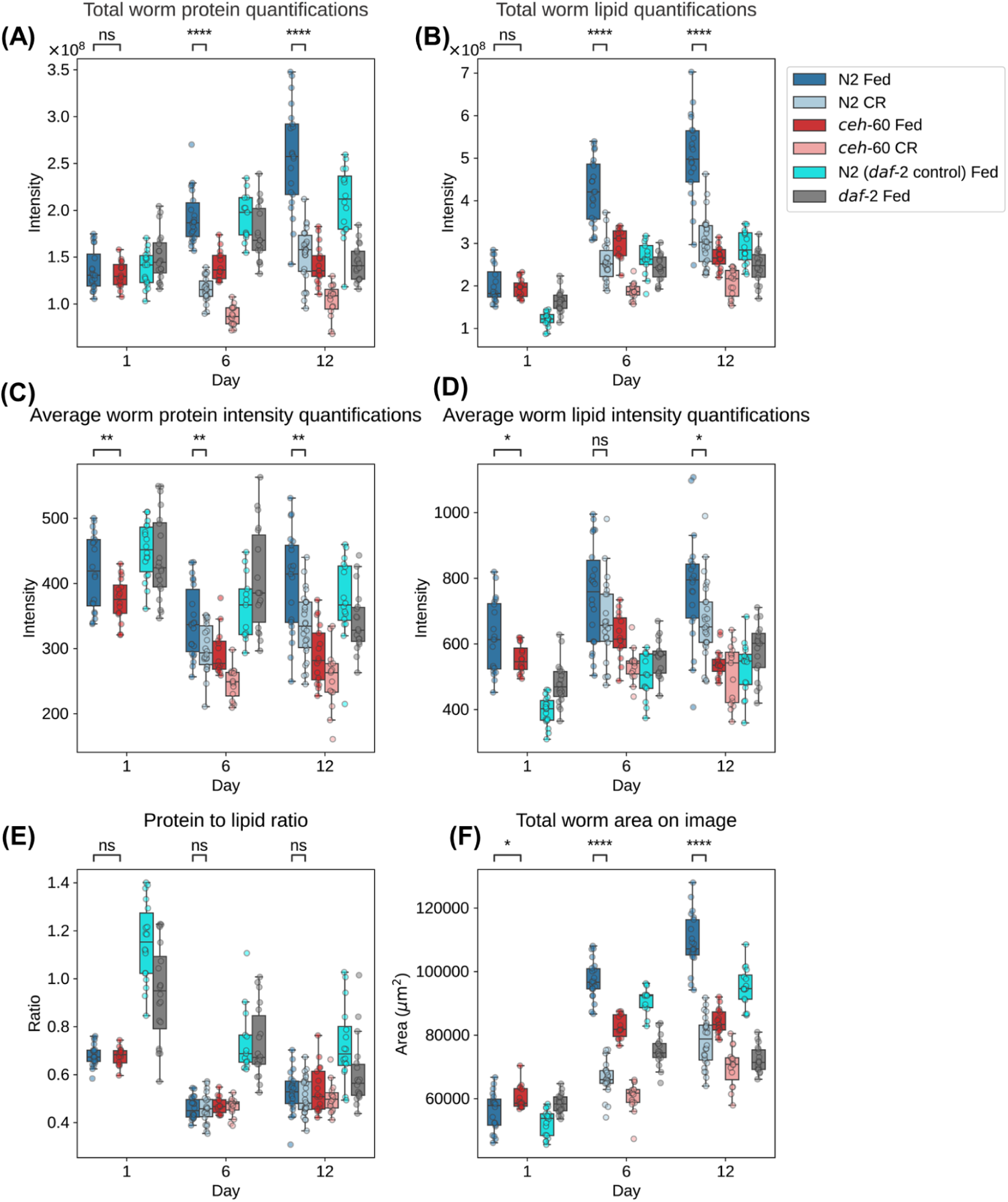
Additional quantification of SRS images. **(A)** Total protein quantification across all conditions **(B)** Total lipid quantification across all conditions **(C)** Total protein normalized to total worm area across all conditions **(D)** Total lipid normalized to total worm area across all conditions **(E)** Total protein signal/total lipid signal across all conditions **(F)** Total area of the worm across all conditions *, **, ***, and **** indicate p < 0.05, 0.01, 0.001, and 0.0001 (Welch’s t-test), respectively. Horizontal bars of the boxes represent maximum, 0.75, 0.5, 0.25, minimum, respectively. Outliers are determined by whether the point deviates 1.5 times the interquartile range (0.75 quartile - 0.25 quartile), from either quartile.

**Figure S2.**
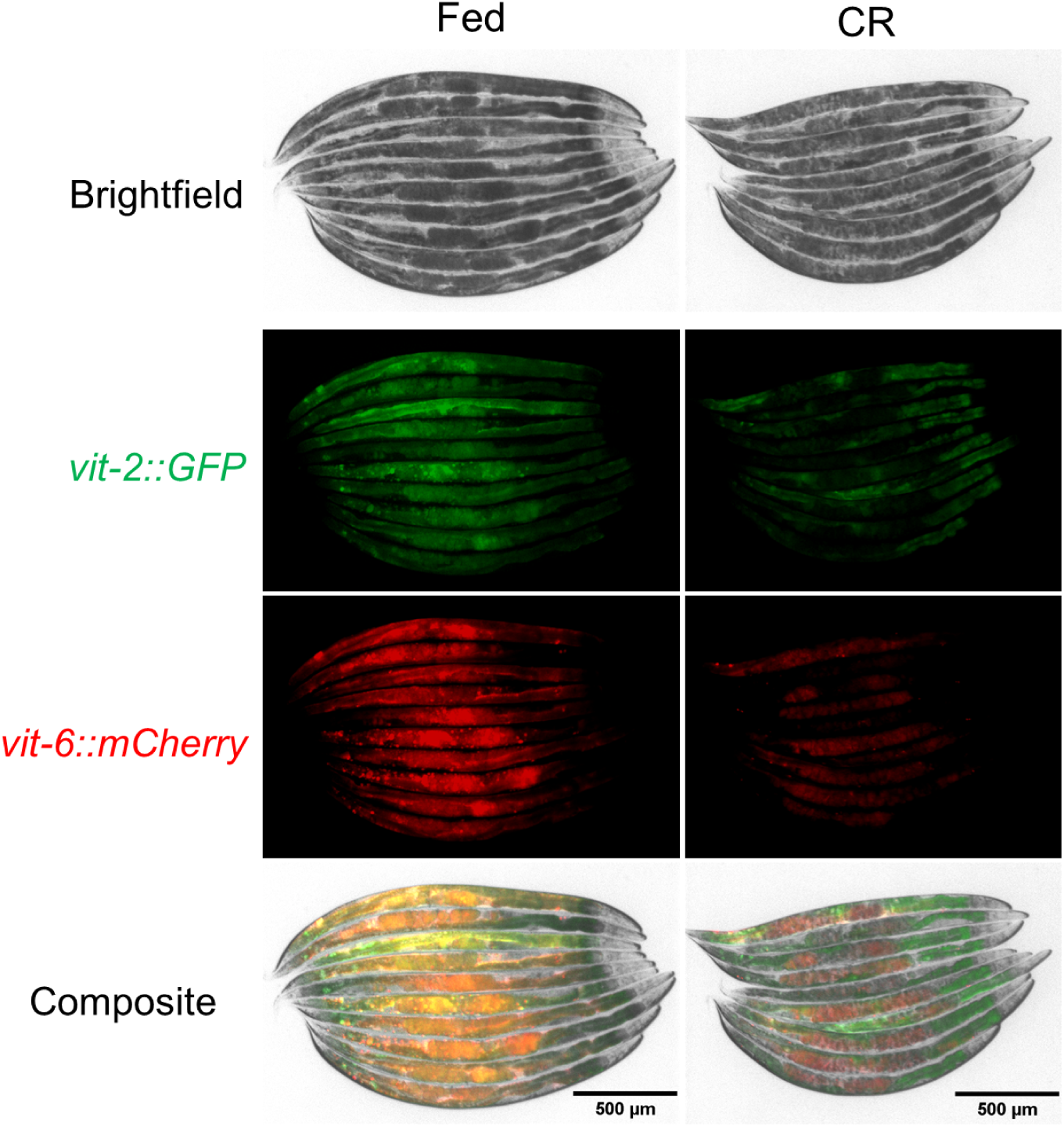
CR decreases vitellogenin protein levels

**Figure S3.**
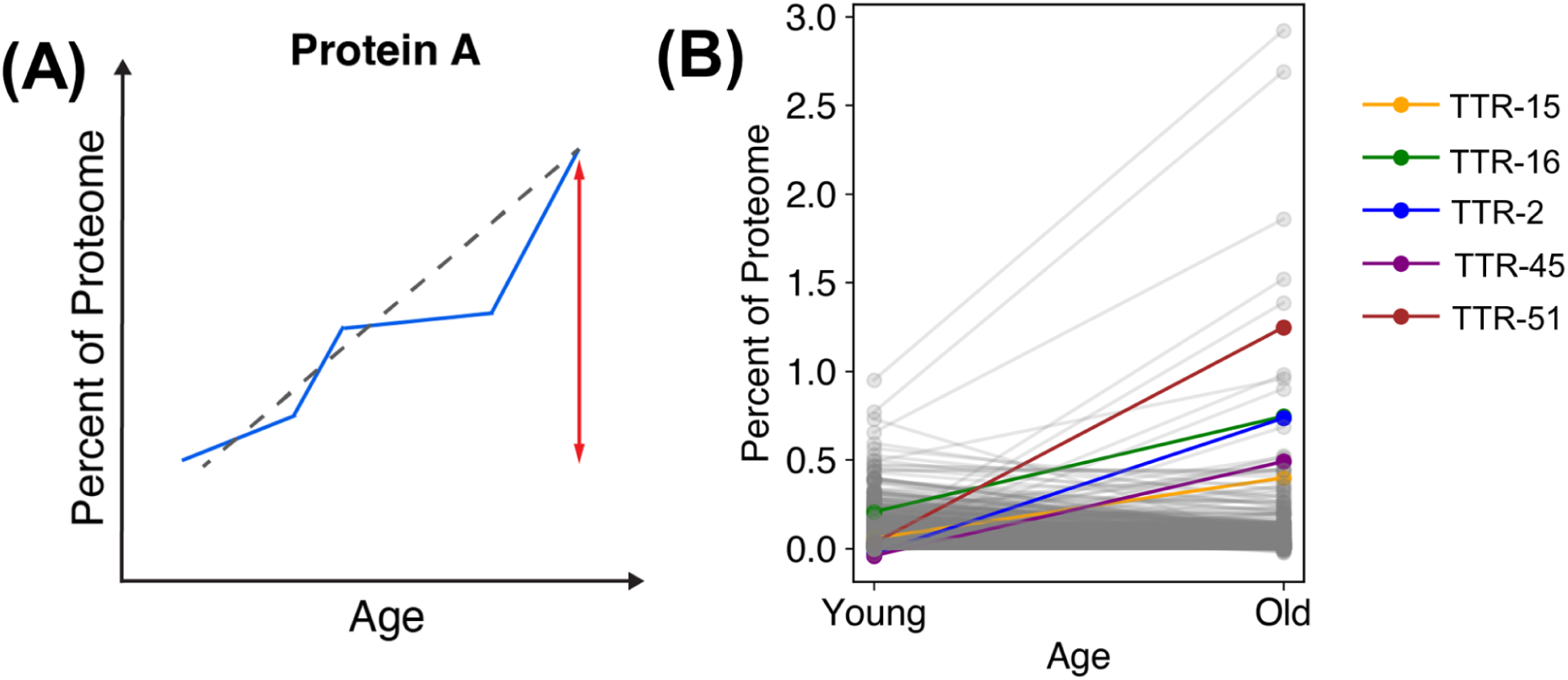
Algorithm for percent proteome calculation and highlighting increase in TTR proteins. **(A)** Schematic of algorithm to calculate largest increase in proteome abundance as fraction of total proteome for each protein identified in proteomics dataset **(B)** TTR proteins increase as a percent of proteome with age

**Figure S4.**
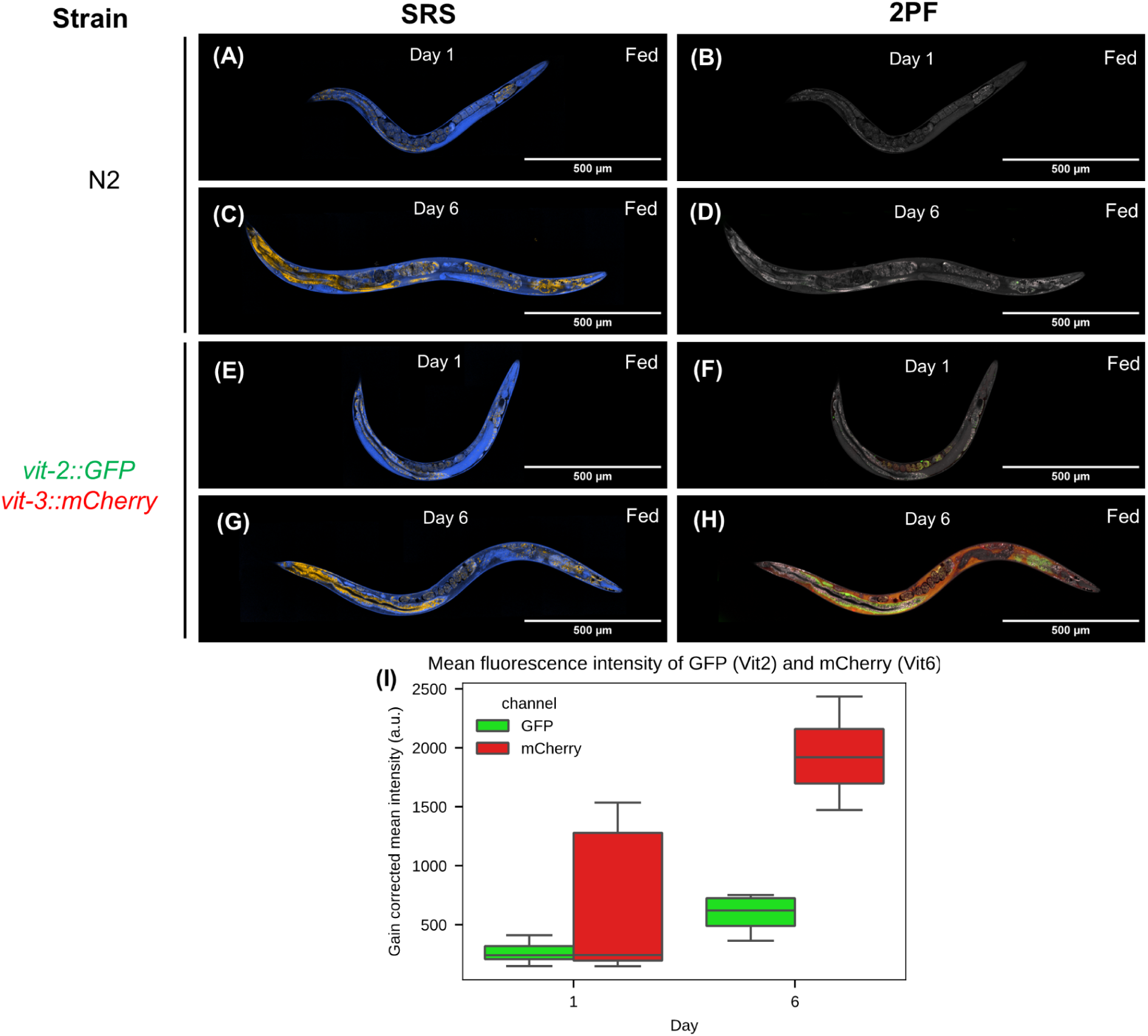
Additional fluorescence control images **(A)** SRS image of Day 1 N2 adult **(B)** Composite SRS and 2PF image of Day 1 N2 adult, SRS CH3 channel in gray, GFP in green, mCherry in red **(C)** SRS image of Day 6 N2 adult **(D)** Composite SRS and 2PF image of Day 6 N2 adult **(E)** SRS image of Day 1 adult tagged with VIT-2::GFP and VIT-3::mcherry **(F)** Composite SRS and 2PF image of Day 1 adult tagged with *vit-2::GFP* and *vit-3::mCherry* **(G)** SRS image of Day 6 adult tagged with VIT-2::GFP and VIT-3::mcherry **(H)** Composite SRS and 2PF image of Day 6 adult tagged with *vit-2::GFP* and *vit-3::mCherry* **(I)** Quantification of VIT-2 and VIT-6 fluorescence intensity with age Horizontal bars of the boxes represent maximum, 0.75, 0.5, 0.25, minimum, respectively. Outliers are determined by whether the point deviates 1.5 times the interquartile range (0.75 quartile - 0.25 quartile), from either quartile.

**Figure S5.**
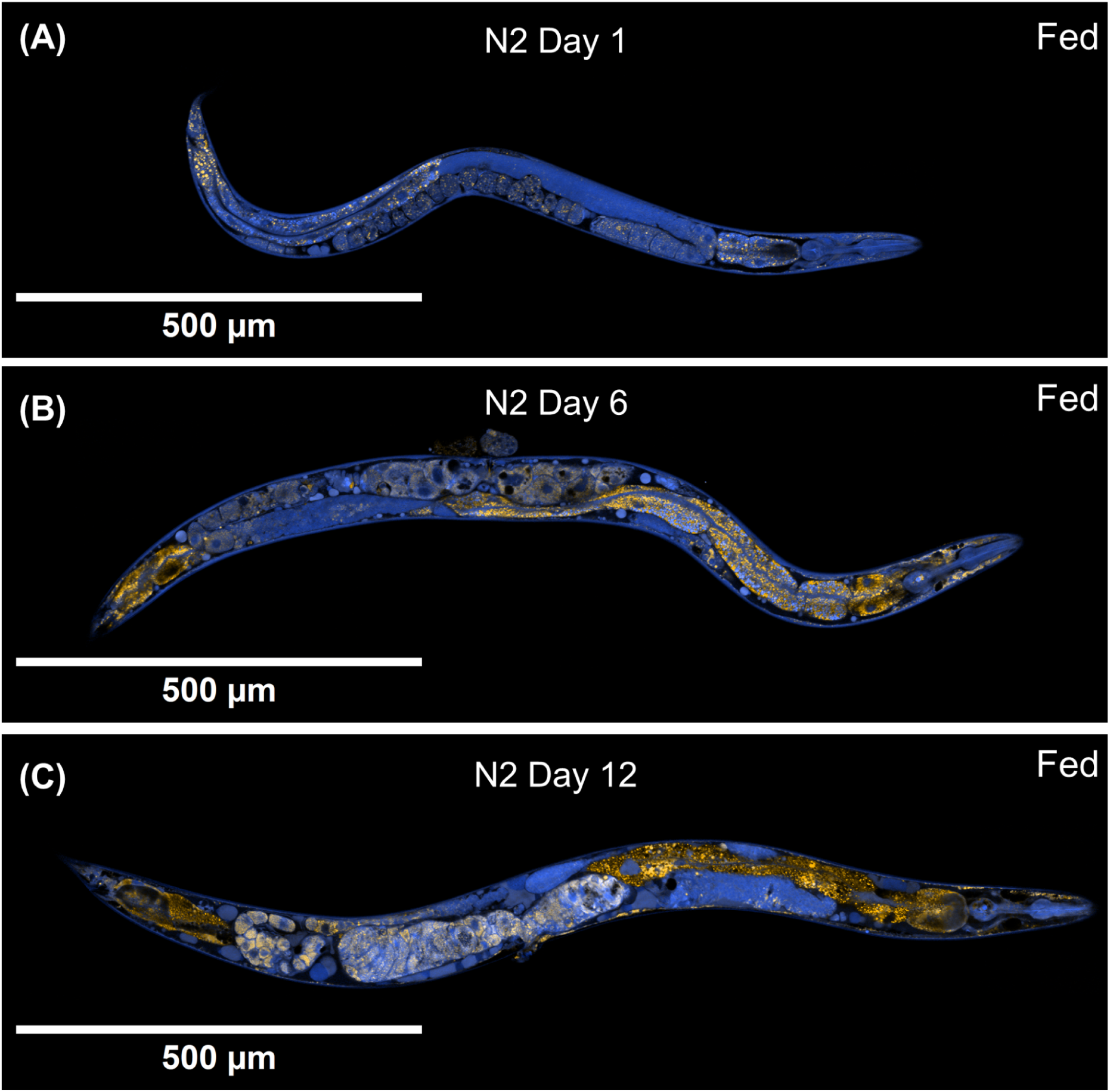
N2 control for *daf-2* worms **(A)** N2 Fed Day 1 adults grown under same conditions as *daf-2* mutants **(B)** N2 Fed Day 6 adults grown under same conditions as *daf-2* mutants **(C)** N2 Fed Day 12 adults grown under same conditions as *daf-2* mutants

**Figure S6.**
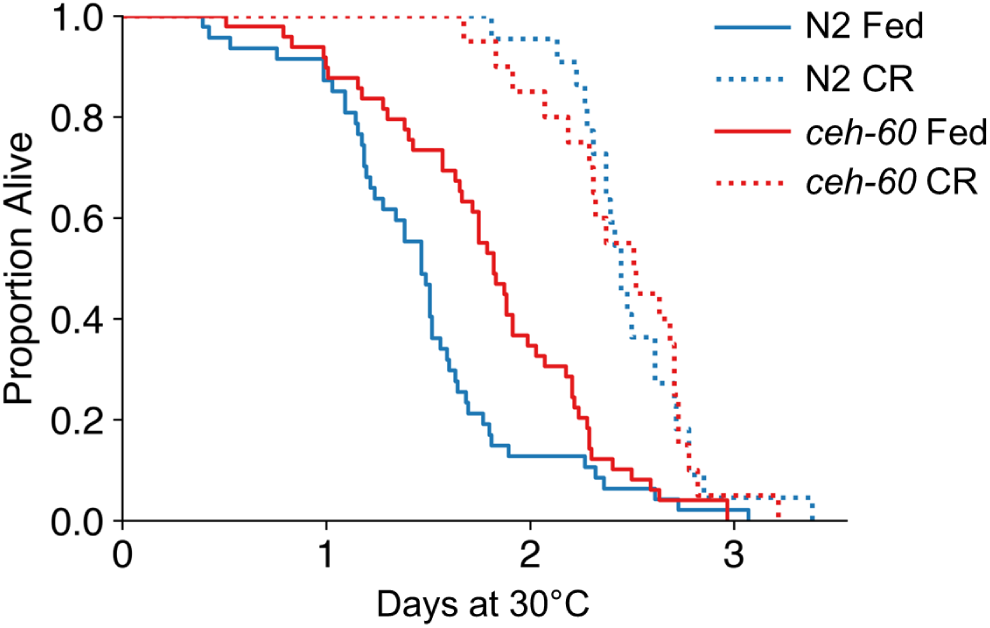
Thermotolerance with N2 and *ceh-60* KO fed and CR

**Figure S7.**
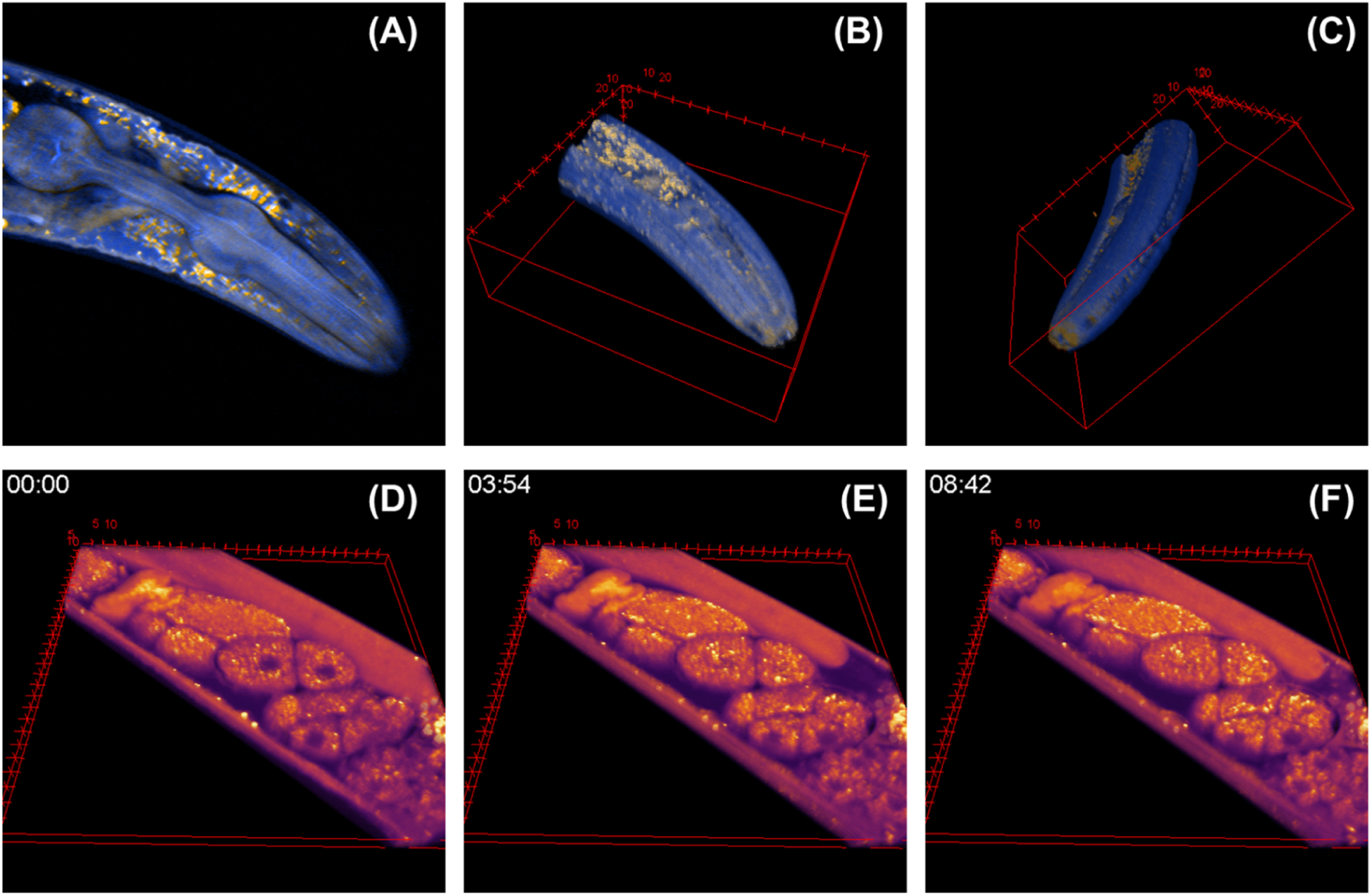
Example of 3D SRS images. **(A)** An SRS image slice from the middle of the worm head imaged in 3D **(B)** 3D rendering of the whole worm head taken in sequential Z-slices **(C)** A rotated view of (B) showing the cuticle of the worm along the side of the head **(D-F)** A timelapse of a small 3D volume in the vicinity of the spermatheca. The gonad of the worm contracts and recovers as the spermatheca fertilizes an oocyte (see attached movie).

## Notes

### Competing Interest Statement

The authors have declared no competing interest.

## REFERENCES

1. López-Otín, C., Blasco, M. A., Partridge, L., Serrano, M. & Kroemer, G. The Hallmarks of Aging. Cell 153, 1194–1217 (2013).

2. Mair, W. & Dillin, A. Aging and Survival: The Genetics of Life Span Extension by Dietary Restriction. Annu. Rev. Biochem. 77, 727–754 (2008).

3. Kenyon, C., Chang, J., Gensch, E., Rudner, A. & Tabtiang, R. A C. elegans mutant that lives twice as long as wild type. Nature 366, 461–464 (1993).

4. Dillin, A. et al. Rates of Behavior and Aging Specified by Mitochondrial Function During Development. Science 298, 2398–2401 (2002).

5. Feng, J., Bussière, F. & Hekimi, S. Mitochondrial Electron Transport Is a Key Determinant of Life Span in Caenorhabditis elegans. Dev. Cell 1, 633–644 (2001).

6. Kimble, J. & Sharrock, W. J. Tissue-specific synthesis of yolk proteins in Caenorhabditis elegans. Dev. Biol. 96, 189–196 (1983).

7. Perez, M. F. & Lehner, B. Vitellogenins - Yolk Gene Function and Regulation in Caenorhabditis elegans. Front. Physiol. 10, 1067 (2019).

8. Sharrock, W. J., Sutherlin, M. E., Leske, K., Cheng, T. K. & Kim, T. Y. Two distinct yolk lipoprotein complexes from Caenorhabditis elegans. J. Biol. Chem. 265, 14422–14431 (1990).

9. Van Nostrand, E. L., Sánchez-Blanco, A., Wu, B., Nguyen, A. & Kim, S. K. Roles of the Developmental Regulator unc-62/Homothorax in Limiting Longevity in Caenorhabditis elegans. PLoS Genet. 9, e1003325 (2013).

10. Ezcurra, M. et al. C. elegans Eats Its Own Intestine to Make Yolk Leading to Multiple Senescent Pathologies. Curr. Biol. 28, 2544–2556.e5 (2018).

11. Murphy, C. T. et al. Genes that act downstream of DAF-16 to influence the lifespan of Caenorhabditis elegans. Nature 424, 277–283 (2003).

12. Seah, N. E. et al. Autophagy-mediated longevity is modulated by lipoprotein biogenesis. Autophagy 12, 261–272 (2016).

13. Dowen, R. H. CEH-60/PBX and UNC-62/MEIS Coordinate a Metabolic Switch that Supports Reproduction in C. elegans. Dev. Cell 49, 235–250.e7 (2019).

14. Mehta, A. & Shapiro, M. D. Apolipoproteins in vascular biology and atherosclerotic disease. Nat. Rev. Cardiol. 19, 168–179 (2022).

15. Baker, M. E. Is vitellogenin an ancestor of apolipoprotein B-100 of human low-density lipoprotein and human lipoprotein lipase? Biochem. J. 255, 1057–1060 (1988).

16. Fontana, L., Meyer, T. E., Klein, S. & Holloszy, J. O. Long-term calorie restriction is highly effective in reducing the risk for atherosclerosis in humans. Proc. Natl. Acad. Sci. 101, 6659–6663 (2004).

17. Borén, J. et al. Low-density lipoproteins cause atherosclerotic cardiovascular disease: pathophysiological, genetic, and therapeutic insights: a consensus statement from the European Atherosclerosis Society Consensus Panel. Eur. Heart J. 41, 2313–2330 (2020).

18. Millar, J. S. et al. Impact of age on the metabolism of VLDL, IDL, and LDL apolipoprotein B-100 in men. J. Lipid Res. 36, 1155–1167 (1995).

19. Freudiger, C. W. et al. Highly specific label-free molecular imaging with spectrally tailored excitation-stimulated Raman scattering (STE-SRS) microscopy. Nat. Photonics 5, 103–109 (2011).

20. Folick, A., Min, W. & Wang, M. C. Label-free imaging of lipid dynamics using Coherent Anti-stokes Raman Scattering (CARS) and Stimulated Raman Scattering (SRS) microscopy. Curr. Opin. Genet. Dev. 21, 585–590 (2011).

21. Wang, M. C., Min, W., Freudiger, C. W., Ruvkun, G. & Xie, X. S. RNAi screening for fat regulatory genes with SRS microscopy. Nat. Methods 8, 135–138 (2011).

22. Wang, P., et al. Imaging Lipid Metabolism in Live Caenorhabditis elegans Using Fingerprint Vibrations. Angew. Chem. Int. Ed. 53, 11787–11792 (2014).

23. Na, H. et al. Identification of lipid droplet structure-like/resident proteins in Caenorhabditis elegans. Biochim. Biophys. Acta BBA - Mol. Cell Res. 1853, 2481–2491 (2015).

24. Mutlu, A. S., Gao, S. M., Zhang, H. & Wang, M. C. Olfactory specificity regulates lipid metabolism through neuroendocrine signaling in Caenorhabditis elegans. Nat. Commun. 11, 1450 (2020).

25. Chen, T., Yavuz, A. & Wang, M. C. Dissecting lipid droplet biology with coherent Raman scattering microscopy. J. Cell Sci. 135, jcs252353 (2021).

26. Fu, D. et al. In Vivo Metabolic Fingerprinting of Neutral Lipids with Hyperspectral Stimulated Raman Scattering Microscopy. J. Am. Chem. Soc. 136, 8820–8828 (2014).

27. Lee, H. J. et al. Assessing Cholesterol Storage in Live Cells and C. elegans by Stimulated Raman Scattering Imaging of Phenyl-Diyne Cholesterol. Sci. Rep. 5, 7930 (2015).

28. Chen, A. J. et al. Fingerprint Stimulated Raman Scattering Imaging Reveals Retinoid Coupling Lipid Metabolism and Survival. ChemPhysChem 19, 2500–2506 (2018).

29. Zhuge, M. et al. Ultrasensitive Vibrational Imaging of Retinoids by Visible Preresonance Stimulated Raman Scattering Microscopy. Adv. Sci. 8, 2003136 (2021).

30. Lin, C.-C. J. & Wang, M. C. Microbial metabolites regulate host lipid metabolism through NR5A–Hedgehog signalling. Nat. Cell Biol. 19, 550–557 (2017).

31. Li, Y., Zhang, W., A. Fung, A. & Shi, L. DO-SRS imaging of metabolic dynamics in aging Drosophila. Analyst 146, 7510–7519 (2021).

32. Li, Y. et al. Direct Imaging of Lipid Metabolic Changes in Drosophila Ovary During Aging Using DO-SRS Microscopy. *Front*. Aging 2, (2022).

33. Li, Y., Zhang, W., Fung, A. A. & Shi, L. DO-SRS imaging of diet regulated metabolic activities in Drosophila during aging processes. Aging Cell 21, e13586 (2022).

34. Mullaney, B. C. & Ashrafi, K. C. elegans fat storage and metabolic regulation. Biochim. Biophys. Acta BBA - Mol. Cell Biol. Lipids 1791, 474–478 (2009).

35. Hou, N. S. & Taubert, S. Function and Regulation of Lipid Biology in Caenorhabditis elegans Aging. Front. Physiol. 3, (2012).

36. Ashrafi, K. et al. Genome-wide RNAi analysis of Caenorhabditis elegans fat regulatory genes. Nature 421, 268–272 (2003).

37. Wang, H. et al. A parthenogenetic quasi-program causes teratoma-like tumors during aging in wild-type C. elegans. Npj Aging Mech. Dis. 4, 6 (2018).

38. Walther, D. M. et al. Widespread Proteome Remodeling and Aggregation in Aging C. elegans. Cell 161, 919–932 (2015).

39. Kimura, K. D., Tissenbaum, H. A., Liu, Y. & Ruvkun, G. *daf-2*, an Insulin Receptor-Like Gene That Regulates Longevity and Diapause in *Caenorhabditis elegans*. Science 277, 942–946 (1997).

40. Zhang, Y.-P. et al. Intestine-specific removal of DAF-2 nearly doubles lifespan in Caenorhabditis elegans with little fitness cost. Nat. Commun. 13, 6339(2022).

41. Wang, M. C., O’Rourke, E. J. & Ruvkun, G. Fat Metabolism Links Germline Stem Cells and Longevity in *C. elegans*. Science 322, 957–960 (2008).

42. Mutlu, A. S., Chen, T., Deng, D. & Wang, M. C. Label-Free Imaging of Lipid Storage Dynamics in Caenorhabditis elegans using Stimulated Raman Scattering Microscopy. J. Vis. Exp. 61870 (2021) doi:10.3791/61870-v.

43. Koyuncu, S. et al. Rewiring of the ubiquitinated proteome determines ageing in C. elegans. Nature 596, 285–290 (2021).

44. Kern, C. C. et al. C. elegans feed yolk to their young in a form of primitive lactation. Nat. Commun. 12, 5801 (2021).

45. DePina, A. S. et al. Regulation of Caenorhabditis elegans vitellogenesis by DAF-2/IIS through separable transcriptional and posttranscriptional mechanisms. BMC Physiol. 11, 11 (2011).

46. Van De Walle, P. et al. CEH-60/PBX regulates vitellogenesis and cuticle permeability through intestinal interaction with UNC-62/MEIS in Caenorhabditis elegans. PLOS Biol. 17, e3000499 (2019).

47. Van Rompay, L., Borghgraef, C., Beets, I., Caers, J. & Temmerman, L. New genetic regulators question relevance of abundant yolk protein production in C. elegans. Sci. Rep. 5, 16381 (2015).

48. Dowen, R. H., Breen, P. C., Tullius, T., Conery, A. L. & Ruvkun, G. A microRNA program in the *C. elegans* hypodermis couples to intestinal mTORC2/PQM-1 signaling to modulate fat transport. Genes Dev. 30, 1515–1528 (2016).

49. Heintz, C. et al. Splicing factor 1 modulates dietary restriction and TORC1 pathway longevity in C. elegans. Nature 541, 102–106 (2017).

50. Panowski, S. H., Wolff, S., Aguilaniu, H., Durieux, J. & Dillin, A. PHA-4/Foxa mediates diet-restriction-induced longevity of C. elegans. Nature 447, 550–555 (2007).

51. Yuan, Y. et al. Enhanced Energy Metabolism Contributes to the Extended Life Span of Calorie-restricted Caenorhabditis elegans. J. Biol. Chem. 287, 31414–31426 (2012).

52. Kapahi, P., Kaeberlein, M. & Hansen, M. Dietary restriction and lifespan: Lessons from invertebrate models. Ageing Res. Rev. 39, 3–14 (2017).

53. Kenyon, C. J. The genetics of ageing. Nature 464, 504–512 (2010).

54. Taylor, R. C. & Dillin, A. Aging as an Event of Proteostasis Collapse. Cold Spring Harb. Perspect. Biol. 3, a004440–a004440 (2011).

55. Douglas, P. M. & Dillin, A. Protein homeostasis and aging in neurodegeneration. J. Cell Biol. 190, 719–729 (2010).

56. Vilchez, D., Saez, I. & Dillin, A. The role of protein clearance mechanisms in organismal ageing and age-related diseases. Nat. Commun. 5, 5659 (2014).

57. Li, X. et al. Quantitative Imaging of Lipid Synthesis and Lipolysis Dynamics in Caenorhabditis elegans by Stimulated Raman Scattering Microscopy. Anal. Chem. 91, 2279–2287 (2019).

58. Zhou, G. et al. Methionine increases yolk production to offset the negative effect of caloric restriction on reproduction without affecting longevity in C. elegans. Aging 12, 2680–2697 (2020).

59. Perez, M. F., Francesconi, M., Hidalgo-Carcedo, C. & Lehner, B. Maternal age generates phenotypic variation in Caenorhabditis elegans. Nature 552, 106–109 (2017).

60. Zhai, C. et al. Fusion and expansion of vitellogenin vesicles during *Caenorhabditis elegans* intestinal senescence. Aging Cell 21, e13719 (2022).

61. Sornda, T. et al. Production of YP170 Vitellogenins Promotes Intestinal Senescence in Caenorhabditis elegans. J. Gerontol. Ser. A 74, 1180–1188 (2019).

62. Blagosklonny, M. V. Aging and Immortality: Quasi-Programmed Senescence and Its Pharmacologic Inhibition. Cell Cycle 5, 2087–2102 (2006).

63. Williams, G. C. PLEIOTROPY, NATURAL SELECTION, AND THE EVOLUTION OF SENESCENCE. Evolution 11, 398–411 (1957).

64. De La Guardia, Y. et al. Run-on of germline apoptosis promotes gonad senescence in *C. elegans*. Oncotarget 7, 39082–39096 (2016).

65. Chotard, L., Skorobogata, O., Sylvain, M.-A., Shrivastava, S. & Rocheleau, C. E. TBC-2 Is Required for Embryonic Yolk Protein Storage and Larval Survival during L1 Diapause in Caenorhabditis elegans. PLoS ONE 5, e15662 (2010).

66. Geens, E. et al. Yolk-deprived *Caenorhabditis elegans* secure brood size at the expense of competitive fitness. Life Sci. Alliance 6, e202201675 (2023).

67. Brawand, D., Wahli, W. & Kaessmann, H. Loss of Egg Yolk Genes in Mammals and the Origin of Lactation and Placentation. PLoS Biol. 6, e63 (2008).

68. Ching, T.-T. & Hsu, A.-L. Solid Plate-based Dietary Restriction in Caenorhabditis elegans. J. Vis. Exp. 2701 (2011) doi:10.3791/2701.

69. Stroustrup, N. et al. The Caenorhabditis elegans Lifespan Machine. Nat. Methods 10, 665–670 (2013).

70. Potma, E. O. Foundations of Nonlinear Optical Microsocpy. (Wiley, 2024).

71. Manifold, B. & Fu, D. Quantitative Stimulated Raman Scattering Microscopy: Promises and Pitfalls. Annu. Rev. Anal. Chem. 15, 269–289 (2022).

72. Manifold, B., Figueroa, B. & Fu, D. Chapter 4 - Hyperspectral SRS imaging via spectral focusing. in *Stimulated Raman Scattering Microscopy* (eds. Cheng, J.-X., Min, W., Ozeki, Y. & Polli, D.) 69–79 (Elsevier, 2022). 10.1016/B978-0-323-85158-9.00035-X.

73. 73. Manifold, B., Dorlhiac, G. F., Landry, M. P. & Streets, A. Imaging neurotransmitter transport in live cells with stimulated Raman scattering microscopy. Preprint at 10.48550/arXiv.2205.05798 (2024).

74. Zhang, X., Dorlhiac, G., Landry, M. P. & Streets, A. Phototoxic effects of nonlinear optical microscopy on cell cycle, oxidative states, and gene expression. Sci. Rep. 12, 18796 (2022).

75. Preibisch, S., Saalfeld, S. & Tomancak, P. Globally optimal stitching of tiled 3D microscopic image acquisitions. Bioinforma. Oxf. Engl. 25, 1463–1465 (2009).

76. Walt, S. van der et al. scikit-image: image processing in Python. PeerJ 2, e453 (2014).

77. Bradski, G. The OpenCV Library. Dr Dobbs J. Softw. Tools (2000).

